# “Integrative Network Meta-Analysis Reveals Estrogen-Mediated RUNX2–PDLIM3–microRNA Crosstalk via ERG Signaling: Implications for Bone and Tissue Regeneration”

**DOI:** 10.1101/2025.11.18.688656

**Authors:** Alshymaa Yusef Hassan

## Abstract

Estrogens govern the female reproductive cycle indefinitely. Estrogens, including estrone (E1), estradiol (E2), estriol (E3), and estetrol (E4), regulate the female life cycle since early embryonic stages and play a crucial role in development, metabolism, and cell function. Throughout evolution, estrogen has regulated reproduction by affecting reproductive organ development and behavior. Estrogen impacts all vertebrates, including fish, and has a role in physiological and pathological states in both genders. The RUNX-2 gene is a member of the RUNX family of transcription factors and encodes a nuclear protein with a Runt DNA-binding domain. This protein is essential for osteoblastic differentiation and skeletal morphogenesis and acts as a scaffold for nucleic acids and regulatory factors involved in skeletal gene expression. The protein can bind DNA both as a monomer or, with more affinity, as a subunit of a heterodimeric complex. In 2022, a study was conducted to characterize novel genes that are regulated by estrogen binding to its receptors (α or β). The PDLIM3 gene, with a coefficient of variation (CV) of 0.083, received the most stable CV score among other genes. Our integrative research uncovers a unique regulatory cascade in which estrogen binding to ERα/β enhances *PDLIM3* expression, then modulating the expression of miR-9, miR-10, and the newly identified miR-6769b, finally activating *RUNX2* transcription.

## I. Introduction

Estrogen, a steroid hormone playing a fundamental role in the female reproductive tract, is an essential regulator of reproductive physiology. Besides its traditional roles, estrogen possesses neuroprotective activity in preventing neurodegenerative disorders such as dementia and reducing the severity of traumatic brain injury. It is also widely utilized in hormone replacement therapy (HRT) for the treatment of symptoms of menstrual irregularities and menopause [1][2]. Among all of the endogenous estrogens, 17β-estradiol (E2) is the most potent and biologically active form found in systemic circulation. E2 regulates a wide array of physiological events in diverse tissues and organs by diffusing through the plasma membrane of target cells and binding to intracellular estrogen receptors (ERs), i.e., ERα and ERβ [2][3]. These bindings trigger cascades of signals that can be divided into genomic and non-genomic mechanisms. In genomic mechanisms, the estradiol-ER complex undergoes a hormone-binding conformational change, translocates into the nucleus, and binds to estrogen response elements (EREs) in enhancer regions, promoters, or untranslated regions of estrogen-responsive genes [4][5]. This receptor-DNA interaction controls gene transcription and subsequent protein synthesis. Otherwise, estrogen may act with membrane-bound receptors such as GPER1 or cytoplasmic ERs in non-genomic signaling, triggering rapid activation of intracellular signaling cascades independent of genomic interaction [6][7]. Both modes of action point to the sophistication and flexibility of the hormone’s function in human physiology. The complex moves to the nucleus and attaches to chromatin at ERE sequences, enhancer regions near promoters, and 30-untranslated regions of target genes. (Figure 1). The RUNX2 gene in humans encodes the transcription factor known as Runt-related transcription factor 2 (RUNX2) or core-binding factor subunit alpha-1 (CBFα1). RUNX2 is recognized as an early marker of osteogenic differentiation and plays a pivotal role in initiating osteoblast-specific extracellular matrix (ECM) synthesis by regulating the expression of critical matrix proteins such as collagen type I and osteopontin (OPN) [8]. It functions as a key transcriptional regulator of osteoblast lineage commitment. RUNX2 encodes a nuclear-localized transcription factor containing a conserved Runt homology domain, which is essential for osteoblast differentiation and skeletal morphogenesis. It operates as a molecular scaffold for nucleic acids and transcriptional co-regulators involved in the control of skeletal gene expression. [9]

**Figure 1.**
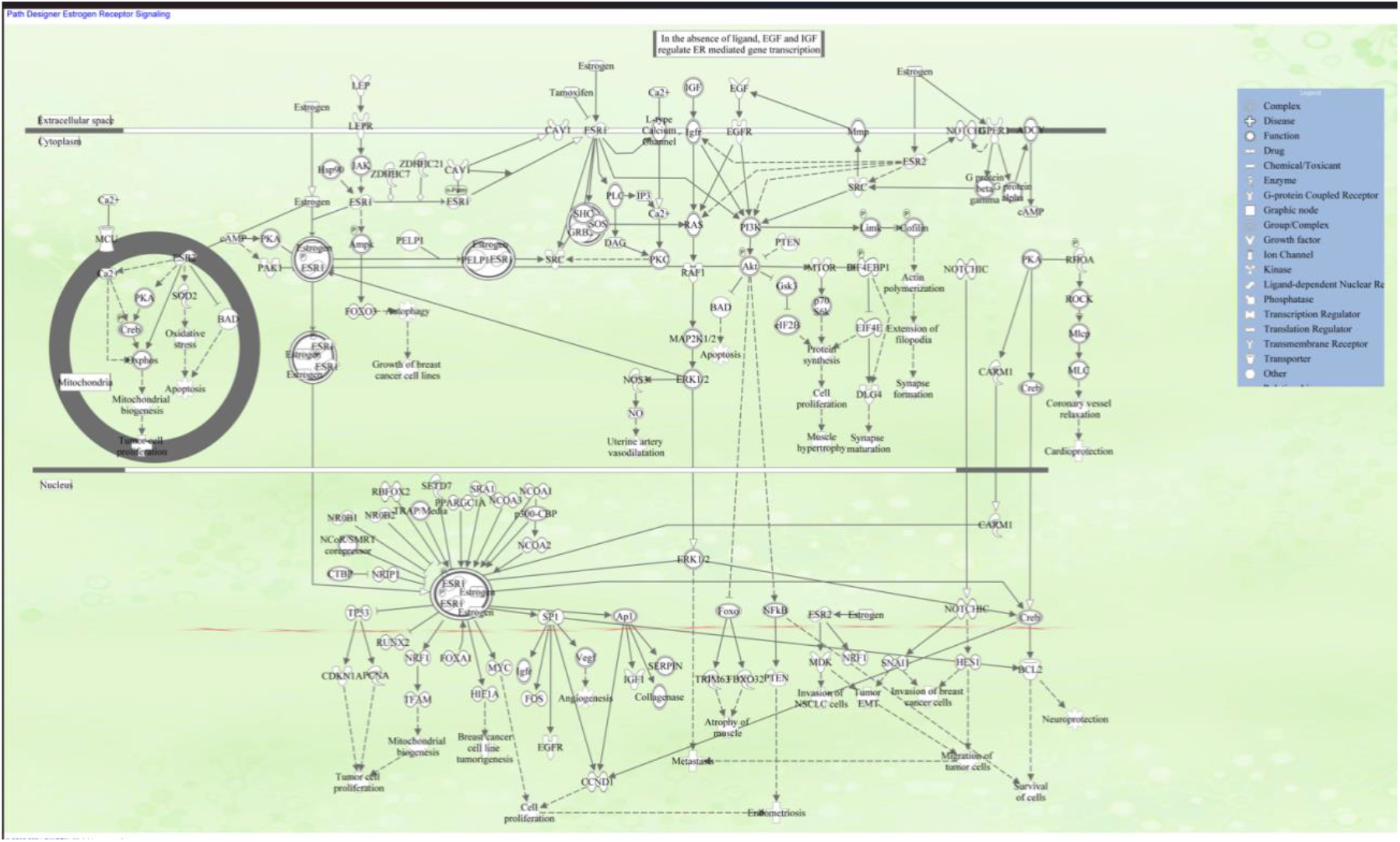
Estrogen receptor signaling, Schematic created using QIAGEN’s Ingenuity Pathway Analysis (IPA) Estrogen receptor signaling pathway, illustrating the key interactions and regulatory mechanisms involved. It includes elements like estrogen receptors (ERα and ERβ), heat shock proteins (HSPs), and various signaling molecules, highlighting processes such as gene expression, cell proliferation, and apoptosis. This pathway plays a crucial role in physiological functions and is particularly important in understanding conditions like breast cancer and hormone-related disorders.ERα, ERβ – Estrogen Receptor Alpha and Beta | E2 – Estradiol | HSP – Heat Shock Protein | SRC – Steroid Receptor Coactivator | PI3K – Phosphoinositide 3-Kinase | AKT – Protein Kinase B | MAPK – Mitogen-Activated Protein Kinase | NF-κB – Nuclear Factor Kappa B | p53 – Tumor Protein p53 | AP-1 – Activator Protein 1 | CREB – cAMP Response Element-Binding Protein | c-Myc – Cellular Myc Protein | STAT – Signal Transducer and Activator of Transcription | EGFR – Epidermal Growth Factor Receptor | GR – Glucocorticoid Receptor | IGF-1 – Insulin-Like Growth Factor 1 | Ras – Small GTPase involved in cell signaling | JNK – c-Jun N-terminal Kinase | Fos – Proto-oncogene c-Fos | Jun – Proto-oncogene c-Jun | VEGF – Vascular Endothelial Growth Factor | Cyclin D – Cell Cycle Regulatory Protein | CDK – Cyclin-Dependent Kinase | PTEN – Phosphatase and Tensin Homolog | GSK3β – Glycogen Synthase Kinase 3 Beta | IRS-1 – Insulin Receptor Substrate 1 | MEK – Mitogen-Activated Protein Kinase Kinase | mTOR – Mechanistic Target of Rapamycin | BCL-2 – B-Cell Lymphoma 2 | BAX – Bcl-2-Associated X Protein---designed using IPA_QIAGEN.

Also, RUNX2 can bind DNA either as a monomer or with greater affinity when part of a heterodimeric complex. The N-terminal domain of the protein includes two potential trinucleotide repeat expansions, which, along with other mutations in the gene, have been implicated in the skeletal disorder cleidocranial dysplasia (CCD) [10].More recently, somatic mutations in the RUNX2 gene, along with its distinct expression signatures in both healthy and neoplastic tissues, have highlighted its prognostic and diagnostic value in multiple human malignancies, supporting its consideration as a cancer biomarker. Studies have demonstrated that RUNX2 contributes to the regulation of essential oncogenic processes, including tumor cell proliferation, angiogenesis, metastasis, cancer stem cell maintenance, and resistance to chemotherapy. These findings underscore the need for deeper investigation into RUNX2-mediated mechanisms as a foundation for novel therapeutic strategies [11].The actin-associated LIM protein (ALP), a product of the PDLIM3 gene also called PDZ and LIM domain protein 3, is a structural and signaling protein found primarily in Z-discs and intercalated discs of cardiac and skeletal muscle tissue. ALP has a key role in the structural integrity of muscle, as it has a role in crosslinking actin filaments via alpha-actinin-2 and is also involved in right ventricle development and functional contractility [12]. Its dysfunction has been implicated in the development of dilated cardiomyopathy (DCM), muscular dystrophy, and tumor growth, highlighting its function in biological and clinical conditions. Despite mounting evidence for the involvement of PDLIM3 in muscle and cardiac physiology, its direct prognostic significance and immunological role within the tumor microenvironment, as in gastric cancer, remain ill-defined [13]. Unexpectedly, in 2022, PDLIM3 was reported as part of a panel of estrogen-responsive genes (ERGs). Estrogen has a considerable effect on gene expression by its interaction with nuclear receptors ERα and ERβ, thus either promoting or suppressing transcriptional activity. Notably, PDLIM3 has been recognized as one of the most sensitive targets under this regulatory mechanism, with a coefficient of variation (CV) of 0.083, which reflects tight regulation by estrogenic signaling [14]. Meanwhile, microRNAs (miRNAs), small non-coding RNAs approximately 19 to 25 nucleotides long, have emerged as key regulators of post-transcriptional gene expression in a variety of developmental and disease settings. Previously disregarded as genomic “noise,” miRNAs are now known to inhibit gene expression, regulate cellular homeostasis, and coordinate responses in diseases from autoimmune disorders to cancer development and viral infection [15].

In this regulatory framework, miR-9 and miR-10 are interesting because of their roles in osteogenic differentiation, whereas mechanistic pathways are still incompletely understood. Western blot studies have shown both miRNAs to influence the expression of Runt-related transcription factor-2 (RUNX2) and the extracellular signal-regulated kinase (ERK) pathway, hypothesizing an intimate interaction between miRNA signaling and osteogenesis [16]. In addition, downregulation of miR-9 in postmitotic neurons is linked to neurodegenerative disorders, emphasizing its role in neuronal survival and maintenance [17]. miR-10 has been demonstrated to suppress T-cell proliferation, induce apoptosis, and facilitate tumor development through multiple models [17]. Interestingly, miR-9 has also been shown to promote differentiation and immunosuppressive activity of myeloid-derived suppressor cells (MDSCs) through targeting Runx1, with possible implications in immunomodulation and tumor immunity [18].

Moreover, miR-10a, a strongly conserved microRNA, has also been involved in various pathological processes, such as rheumatoid arthritis [19], juvenile dermatomyositis [20], and a range of cancers [21], underlining its therapeutic and diagnostic utility in a variety of clinical settings.Previous functional studies have also corroborated the idea that miR-10a-3p actively suppresses the production of Inhibitors of Differentiation (ID) genes ID3, boosting the activity of the ossification core factor RUNX2. [22] Another microRNA we would like to highlight is microRNA6769B (mammalian), which was discovered to have an indirect regulatory influence on the expression of the RUNX-2 gene via miR-1896 (and other miRNAs w/seed GGUGGGU) (mammalian) activation, leading to upregulation of downstream signaling pathways.

The purpose of our research is to shed light on current mechanistic findings and the modulatory role of estrogen via both direct and indirect effects on the signaling pathways that regulate RUNX-2 expression. Depending on the data analyzed, our primary goal is to link the expression and regulation of microRNA9, microRNA10, miR-1896, and microRNA6769B to RUNX-2 expression by modulating the new estrogen receptor gene, ERG-PDLIM3. QIAGEN’s bioinformatics tool, Ingenuity Pathway Analysis (IPA), was used to design molecular networks and analyze their biological roles. The molecular networks were compared to QIAGEN Knowledge Base (QKB) findings using canonical and signaling pathway analysis, as well as other statistical approaches.

## II. Material and Methods

### Ingenuity Pathway Analysis Software

IPA, a bioinformatics software tool for data mining, uses canonical pathways and gene regulatory networks from literature to help interpret and analyze various biological pathways. Various techniques were used to create pathways depicting the molecular networks connected with estrogen, RUNX-2, different microRNAs, and their intermediary molecules to evaluate functional hypotheses. The bioinformatics tool utilized data from the QIAGEN Knowledge Base (QKB) between February 5th, 2024, and June 14th, 2025. [23] [24] [25][26]. Figure 2 illustrates the workflow utilized from QIAGEN’s Ingenuity Pathway Analysis (IPA) bioinformatics software for data mining.

**Figure 2.**
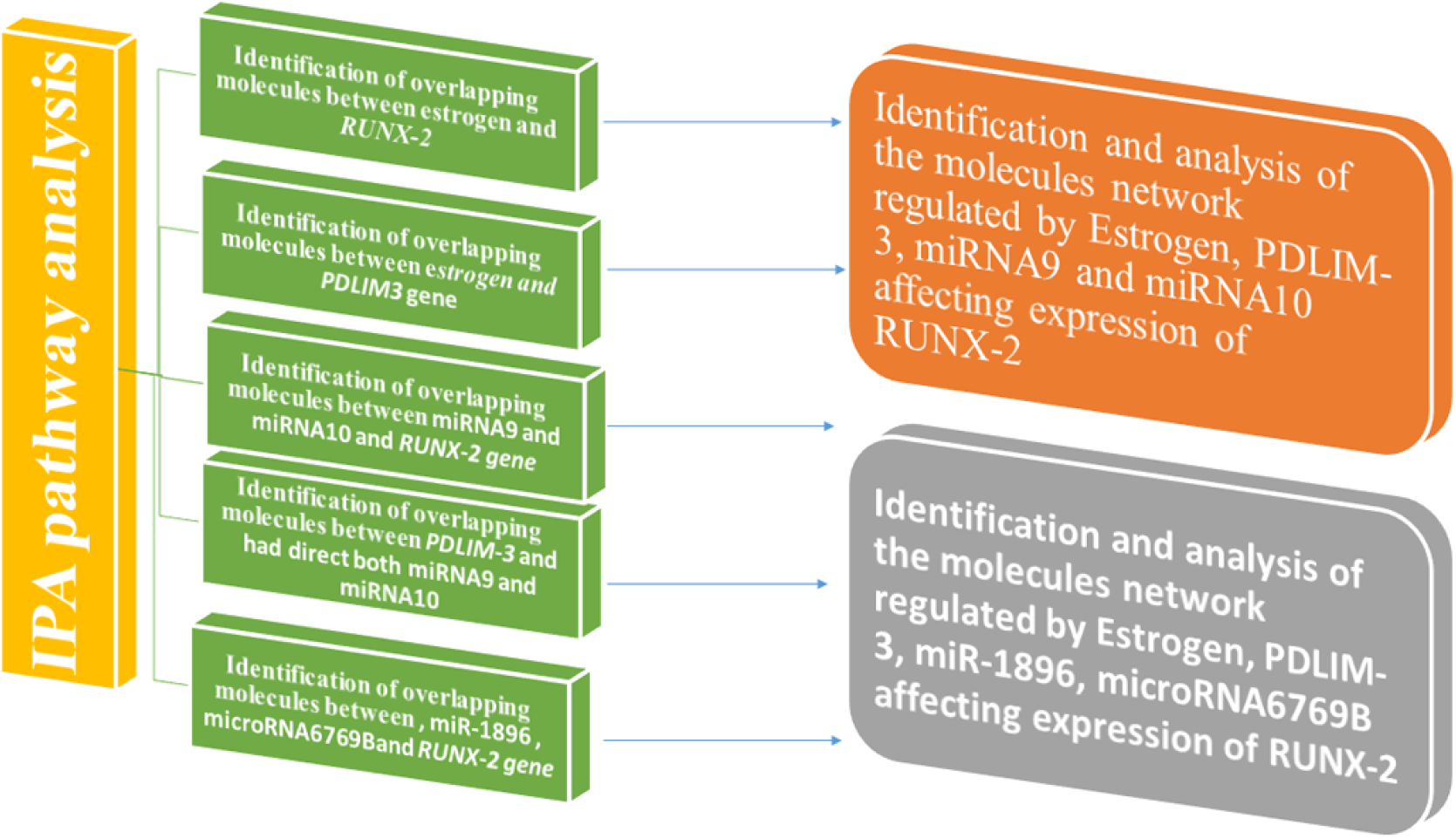
The data mining workflow was based on QIAGEN’s Ingenuity Pathway Analysis (IPA) bioinformatics tools. The “Grow”, “Connect”, “Pathway Explorer”, and “Molecule Activity Predictor” (MAP) tools from the “My Pathway” feature were used to develop biological networks that showed the connectivity between distinct nodes. Furthermore, the “Core Analysis: Expression Analysis” tool was utilized to compare the molecules within the produced molecular route to canonical pathways recorded within QIAGEN’s knowledge base (QKB).

## III. Results and Outcomes, all the figures and tables are provided as supplementary materials

### 1. Molecular Pathway Analysis of Molecules Mediating the Relationship Between Estrogen and *PDLIM3*

The “MAP” program was used to generate a connectivity map depicting the interaction between the 10 molecules involved with estrogen’s direct influence and their relationship with *PDLIM3*. This is depicted in Figure 3 and Supplemental Tables 1–2. Our findings show that estrogen is associated with intermediates such as Proliferating Cellular Nuclear Antigen (PCNA), Cyclin-Dependent Kinase 4 (CDK4), Luteinizing Hormone (LH), Mitogen-Activated Protein Kinase-1 (MAPK-1), Estrogen Receptor 1 and 2 (ESR 1 & 2), and Follicle-Stimulating Hormone (FSH), all of which are linked to the *PDLIM3* gene. This indicates that estrogen’s modulation of PDLIM3 may be closely associated with cell cycle progression and gonadotropin signaling pathways, placing PDLIM3 at the nexus of hormonal and proliferative signals.

### 2. Molecular Pathway Analysis of Molecules Mediating the Relationship Between Estrogen and Affect Expression of RUNX-2

The “MAP” program and QKB can identify 75 estrogen-controlled pathways that regulate RUNX-2 expression. These molecules included biological medicines, the canonical pathway, complexes, cytokines, enzymes, G-protein coupled receptors, kinases, nuclear receptors, peptidase, phosphatase, transcription and translation regulators, transmembrane receptors, and transporters, as depicted in Figure 4 and Tables 3 and 4, respectively. The magnitude and variety of this network highlight the pivotal and multifaceted role of estrogen as a principal regulator of *RUNX-2*, able to affect osteogenesis through an extensive array of signaling pathways.

### 3. Molecular Pathway Analysis of Molecules Mediating the Relationship Between miRNA9 and Directly Affecting Expression of the RUNX-2 Gene

The “MAP” program and QKB identified 8 pathways controlled by miRNA9 that modulate RUNX-2 expression. These molecules comprised G-protein coupled receptors, kinases, nuclear receptors, transcription and translation regulators, transmembrane receptors, and transporters (Figure 5 and Tables 5 and 6, respectively). This targeted and substantial network identifies miR-9 as a distinct and powerful epigenetic modulator that precisely regulates the expression of the osteogenic master regulator *RUNX-2*.

### 4. Molecular Pathway Analysis of Molecules Mediating the Relationship Between miRNA10 and Directly Affecting Expression of the RUNX-2 Gene

The “MAP” program and QKB identified 34 pathways controlled by miRNA10 that modulate RUNX-2 expression. These molecules comprised G-protein coupled receptors, kinases, nuclear receptors, transcription and translation regulators, transmembrane receptors, and transporters, as shown in Figure 6 and Tables 7 and 8, respectively. The extensive regulatory scope of miR-10, in contrast to miR-9, implies it may function as a superior integrator, harmonizing several signals to facilitate RUNX-2 activation.

### 5. Molecular Pathway Analysis of Molecules Mediating the Relationship Between *PDLIM-3* and Had Direct Effects on Expression of miRNA9

The “MAP” program and QKB identified 49 pathways directly regulated by *PDLIM-3* that modulate miRNA9 expression. These molecules comprised G-protein coupled receptors, kinases, nuclear receptors, transcription and translation regulators, transmembrane receptors, and transporters, as shown in Figure 7 and Tables 9 and 10, respectively. This discovery establishes a molecular connection, demonstrating how PDLIM3, a structural protein, functions as a signaling hub to epigenetically modulate gene expression by regulating miR-9 levels.

### 6. Molecular Pathway Analysis of Molecules Mediating the Relationship Between *PDLIM-3* and Had Direct Effects on Expression of miRNA10

The “MAP” program and QKB identified 98 pathways directly regulated by *PDLIM-3* that modulate miRNA9 expression. These molecules comprised G-protein coupled receptors, kinases, nuclear receptors, transcription and translation regulators, transmembrane receptors, and transporters, as shown in Figure 8 and Tables 11 and 12, respectively. This is the most comprehensive connectivity in our data, strongly implying that the regulation of miR-10 is a significant downstream function of PDLIM3, probably its principal role in the estrogen-mediated signaling pathway leading to osteogenesis.

### 7. Identification and Analysis of the Molecule Network Regulated by Estrogen, PDLIM-3, miRNA9, and miRNA10 affects the expression of RUNX-2

The “MAP” program and QKB identified around 39 molecules involved directly or indirectly in the expression of *RUNX-2*, regulated by estrogen, miRNA9, miRNA10, and *PDLIM-3* interaction.

The full classification of those molecules, including the gene name, its location, and its family, is shown in Figure 9 and Tables 13 and 14, respectively. This integrated map consolidates our findings, visually delineating the novel multi-layered regulatory pathway from estrogen receptor stimulation to the transcriptional regulation of RUNX-2.

### 8. Identification and Analysis of the Molecules Network Regulated by Estrogen, PDLIM-3, miR-1896, microRNA6769B affects the expression of RUNX-2

The “MAP” program and QKB identified around 45 molecules involved directly or indirectly in the expression of RUNX-2, regulated by estrogen, microRNA-1896, microRNA-6769B, and *PDLIM-3*. The full classification of those molecules, including the gene name, its location, and its family, is shown in Figure 10 and Tables 15 and 16, respectively. This not only confirms the function of PDLIM3 as an epigenetic regulator but also considerably broadens the suggested regulatory axis by incorporating miR-6769b as a novel and possibly crucial participant in estrogen-responsive osteogenesis.

## IV. Constraints

This research utilizes integrative network modeling with the QIAGEN Ingenuity Pathway Analysis (IPA) platform and the QIAGEN Knowledge Base. The results depend on curated datasets, documented interactions, and the completeness of databases, rather than on de novo transcriptome experiments. Despite offering mechanistic insight, the proposed estrogen–PDLIM3–microRNA– RUNX2 regulation axis has not been experimentally confirmed using functional assays, gene editing, or live-cell models. IPA does not establish causality beyond recognized interactions, and its enrichment results are influenced by existing literature bias. Consequently, the suggested signaling model ought to be regarded as hypothesis-generating, necessitating focused experiments—such as CRISPR-mediated disruption, qPCR validation, or estrogen-response assays—to verify the functional relevance of PDLIM3 and the identified microRNAs. Nevertheless, the integrative signal identified across various IPA functions establishes a robust basis for forthcoming mechanistic investigation.

## V. Discussion and Conclusion

This study elucidates a novel, multilayered regulatory axis in which estrogen activates RUNX2 expression through both direct genomic signaling and epigenetic modulation involving PDLIM3 and specific microRNAs. The identification of *PDLIM3* as an intermediary, along with the integration of miR-9, miR-10, and the newly implicated miR-6769b, offers novel perspectives on estrogen-responsive osteogenesis and opens potential avenues for targeted therapeutic strategies.

Estrogen-driven regulatory cascade linking the ERα/β–PDLIM3–miRNAs–RUNX2 axis.

The pathway identified can be summarized as

### 1. Classical Estrogen Receptor Signaling and PDLIM3 Activation

Estrogen binding to ERα and ERβ activates classical genomic signaling, whereby the ligand-receptor complex translocates to the nucleus and binds estrogen response elements (EREs) in the regulatory regions of target genes [27]. Our analysis revealed a strong association between estrogen signaling and *PDLIM3* expression. Although *PDLIM3* has previously been associated primarily with muscle function [28], this is the first report linking it to estrogen-mediated osteogenic pathways and RUNX2 regulation. Its identification as a novel intermediary in this axis is supported by its low coefficient of variation (CV = 0.083), indicating tight estrogenic regulation [29].

### 2. PDLIM3-Mediated Regulation of miR-9 and miR-10

Downstream of *PDLIM3*, our IPA-based analysis identified significant associations with miR-9 and miR-10, two miRNAs implicated in bone formation, neurodevelopment, and immune regulation [30–31]. Both miRNAs have been shown to regulate *RUNX2* directly or via ERK signaling, and their increased expression in response to estrogen-PDLIM3 signaling suggests a crucial post-transcriptional modulatory role in osteogenesis [32].

### 3. Novel Discovery of miR-6769b in Osteogenic Regulation

An especially noteworthy finding is the identification of miR-6769b as a novel epigenetic player within this regulatory framework. While limited prior data exist linking miR-6769b to bone biology, emerging literature suggests its involvement in exosome-mediated bone remodeling and cell proliferation control [33,34]. Our network data suggest that miR-6769b is induced downstream of PDLIM3, representing a new layer of epigenetic regulation for RUNX2.

### 4. Integrated Signaling Cascade

Our findings support a multi-step model for estrogen-driven RUNX2 activation:

Estrogen → ERα/β → PDLIM3 ↑ → miR-9/miR-10/miR-6769b ↑ → RUNX2 ↑

This pathway expands upon the classical model of estrogen action by incorporating miRNA-mediated epigenetic regulation and identifying new targets such as PDLIM3 and miR-6769b that can serve as biomarkers or therapeutic entry points in skeletal disorders [27, 29].

### 5. Implications and Future Directions

1. Biological Significance: This model enriches our understanding of estrogenic regulation of skeletal development and uncovers new candidate targets for bone regeneration and disease treatment.
2. Experimental Validation: Further experimental validation using ChIP-qPCR (to confirm ER binding to PDLIM3), gene knockdown (for PDLIM3 and miR-6769b), and miRNA modulation studies is critical to establish functional causality.

## VI. Conclusion

In this study, pathway relationships between estrogen, *PDLIM3*, microRNAs (miR-9, miR-10, and miR-6769b), and RUNX2 were primarily modeled using QIAGEN’s Ingenuity Pathway Analysis (IPA) tools, in conjunction with curated biological insights from literature. While z-score-based predictions are commonly employed to infer activation or inhibition states in large-scale differential expression studies, they were not the focus of this analysis. Instead, we aimed to systematically map molecular interactions and delineate hierarchical regulatory networks rather than statistically quantify expression changes.

Consequently, the findings are presented descriptively, highlighting directionality and mechanistic connectivity as supported by canonical pathway data and peer-reviewed evidence. This approach has uncovered a novel, multilayered regulatory cascade whereby estrogen signaling through ERα/β leads to upregulation of *PDLIM3*, which in turn enhances the expression of key microRNAs, including miR-9, miR-10, and the newly implicated miR-6769b, ultimately driving the transcriptional activation of RUNX2.

These insights reveal new mediators within the estrogen–RUNX2 axis and offer promising implications for future research into skeletal development, osteogenic differentiation, and hormone-responsive bone pathologies such as osteoporosis and fracture repair.

## Supporting information

https://doi.org/10.5281/zenodo.17089993

## VII. List of Abbreviations

Abbreviation: Definition
AKT: Protein kinase B; serine/threonine kinase in the PI3K/AKT signaling pathway regulating survival and metabolism.
ALP: Actin-associated LIM protein; structural and signaling protein encoded by **PDLIM3**, localized in cardiac and skeletal muscle Z-discs.
AP-1: Activator protein 1; transcription factor complex (Fos/Jun) regulating proliferation and apoptosis.
BAX: Bcl-2–associated X protein; pro-apoptotic member of the Bcl-2 family.
BCL-2: B-cell lymphoma 2; anti-apoptotic protein that promotes cell survival.
CBFα1: Core-binding factor subunit alpha-1; alternative name for RUNX2 transcription factor.
CCD: Cleidocranial dysplasia; hereditary skeletal disorder caused by mutations in **RUNX2**.
CDK: Cyclin-dependent kinase; enzyme family that regulates cell-cycle progression.
CDK4: Cyclin-dependent kinase 4; G1 phase cell-cycle regulator.
CREB: cAMP response element-binding protein; transcription factor that regulates metabolism, survival, and plasticity.
CV: Coefficient of variation; measure of relative variability in gene expression or stability.
DCM: Dilated cardiomyopathy; disorder characterized by dilation and impaired contraction of cardiac chambers.
E1: Estrone; endogenous estrogen.
E2: 17β-estradiol; the most potent and biologically active endogenous estrogen.
E3: Estriol; estrogen predominant during pregnancy.
E4: Estetrol; fetal liver–derived estrogen present during pregnancy.
ECM: Extracellular matrix; structural network of proteins and polysaccharides surrounding cells.
EGFR: Epidermal growth factor receptor; receptor tyrosine kinase regulating proliferation and survival.
ER: Estrogen receptor; ligand-activated transcription factor mediating estrogen actions.
ERα / ERβ: Estrogen receptor alpha / beta; two main nuclear estrogen receptor isoforms.
ERGs: Estrogen-responsive genes; genes whose transcription is modulated by estrogen receptor signaling.
EREs: Estrogen response elements; DNA sequences bound by ER complexes to regulate transcription.
ERK: Extracellular signal-regulated kinase; MAPK family kinase involved in proliferation and differentiation.
ESR1 / ESR2: Estrogen receptor 1 / 2; gene symbols encoding ERα and ERβ, respectively.
FSH: Follicle-stimulating hormone; gonadotropin regulating follicle development and spermatogenesis.
Fos: Proto-oncogene c-Fos; component of the AP-1 transcription factor complex.
GPER1: G protein-coupled estrogen receptor 1; membrane-bound estrogen receptor mediating rapid non-genomic signaling.
GR: Glucocorticoid receptor; nuclear receptor for glucocorticoid hormones.
GSK3β: Glycogen synthase kinase 3 beta; serine/threonine kinase involved in metabolism and Wnt/MAPK signaling.
HRT: Hormone replacement therapy; clinical use of hormones (e.g., estrogens) to treat menopausal or deficiency symptoms.
HSPs: Heat shock proteins; molecular chaperones that stabilize and refold proteins, also associated with steroid receptors.
ID: Inhibitor of differentiation; family of helix-loop-helix transcriptional regulators.
ID3: Inhibitor of DNA-binding protein 3; an ID family member downregulated by miR-10a-3p during osteogenic differentiation.
IGF-1: Insulin-like growth factor 1; peptide growth factor important for growth and metabolism.
IPA: Ingenuity Pathway Analysis; QIAGEN bioinformatics software for pathway and network modeling.
IRS-1: Insulin receptor substrate 1; adaptor protein transmitting insulin/IGF-1 receptor signaling.
JNK: c-Jun N-terminal kinase; stress-activated protein kinase within the MAPK family.
LH: Luteinizing hormone; gonadotropin regulating ovulation and gonadal steroid production.
MAP: Molecule Activity Predictor; Ingenuity Pathway Analysis (IPA) tool predicting activation/inhibition effects within networks.
MAPK / MAPK-1: Mitogen-activated protein kinase / MAPK-1; serine/threonine kinases mediating downstream signaling (ERK2 often referred to as MAPK-1).
MDSCs: Myeloid-derived suppressor cells; immune-suppressive myeloid cell population modulating tumor and inflammatory responses.
MEK: Mitogen-activated protein kinase kinase; dual-specificity kinase upstream of ERK in the MAPK cascade.
miR / miRNA: MicroRNA; small (∼19–25 nt) non-coding RNA that regulates gene expression post-transcriptionally.
miR-9: MicroRNA-9; miRNA regulating neurogenesis, osteogenesis, and immune functions.
miR-10 / miRNA10 / miR-10a: MicroRNA-10; family associated with differentiation, apoptosis, and cancer; miR-10a is a specific isoform.
miR-1896: MicroRNA-1896; miRNA implicated in indirect regulation of RUNX2 via shared seed sequences.
miR-6769b / microRNA6769B: MicroRNA-6769b; newly implicated epigenetic regulator in RUNX2-related signaling.
mTOR: Mechanistic target of rapamycin; central kinase controlling growth and metabolism.
NF-κB: Nuclear factor kappa-B; transcription factor regulating inflammatory and immune responses.
OPN: Osteopontin; extracellular matrix phosphoprotein involved in bone remodeling and mineralization.
PCNA: Proliferating cell nuclear antigen; sliding clamp protein and marker of DNA replication and repair.
PDLIM3: PDZ and LIM domain protein 3; actin-associated structural and signaling protein (ALP), here identified as an estrogen-regulated intermediary upstream of RUNX2.
PI3K: Phosphoinositide 3-kinase; lipid kinase that generates PIP3 and activates AKT signaling.
PTEN: Phosphatase and tensin homolog; tumor suppressor and negative regulator of PI3K/AKT signaling.
QKB: QIAGEN Knowledge Base; curated database underpinning Ingenuity Pathway Analysis.
Ras: Rat sarcoma; small GTP-binding protein family regulating proliferation and survival pathways.
RUNX2 / RUNX-2: Runt-related transcription factor 2; master transcription factor for osteoblast differentiation and skeletal morphogenesis.
SRC: Steroid receptor coactivator; transcriptional co-regulator that enhances nuclear receptor– mediated gene expression.
STAT: Signal transducer and activator of transcription; transcription factor family that mediates cytokine and growth factor signaling.
VEGF: Vascular endothelial growth factor; central regulator of angiogenesis and vascular permeability.

## VIII. Acknowledgments

I would like to express my deepest gratitude to Professor Sulie Chang and the Institute of Neuroimmune Pharmacology (INIP) at Seton Hall University for their continuous support, guidance, and mentorship throughout my graduate studies. Their invaluable contributions and encouragement have played a significant role in the development of this work and my academic growth.

## XI. Figure Legends

a. Figure 1. Diagram of Estrogen Receptor Signaling Pathway. This figure, produced with QIAGEN’s Ingenuity Pathway Analysis (IPA), depicts the principal molecular connections and regulatory mechanisms of estrogen receptor signaling. It encompasses fundamental elements such estrogen receptors (ERα and ERβ), heat shock proteins (HSPs), and downstream signaling molecules, emphasizing mechanisms such as gene expression, cellular proliferation, and apoptosis.
b. Figure 2. Workflow for Bioinformatics Data Mining and Analysis. A schematic depiction of the data analysis process employing QIAGEN’s Ingenuity Pathway Analysis (IPA) technologies. The method illustrates the utilization of the “Grow,” “Connect,” “Pathway Explorer,” and “Molecule Activity Predictor” (MAP) tools for constructing and analyzing biological networks, then comparing them to canonical pathways in the QIAGEN Knowledge Base (QKB).
c. Figure 3. Molecular Network Connecting Estrogen Signaling to PDLIM3 Expression. A connection map produced with the IPA “MAP” tool, illustrating the direct relationships between estrogen and the PDLIM3 gene through 10 principal intermediary molecules, such as PCNA, CDK4, ESR1, and ESR2. This network indicates that estrogen modulates PDLIM3 via pathways associated with cell cycle progression and gonadotropin signaling.
d. Figure 4. Comprehensive Network of Estrogen-Regulated Pathways Modulating RUNX2 Expression. The “MAP” software and QKB found 75 unique molecular pathways by which estrogen regulates RUNX2 expression. This network includes several molecules, such as transcription regulators, kinases, and cytokines, highlighting estrogen’s multifaceted role as a principal regulator of osteogenesis.
e. Figure 5. Regulatory Network of miRNA-9 Targeting RUNX2. The molecular network illustrates eight pathways by which miRNA-9 directly influences RUNX2 expression, encompassing transcription regulators and other molecular families. This designates miR-9 as a precise epigenetic modulator of the osteogenic master regulator.
f. Figure 6. Extensive Regulatory Network of miRNA-10 Targeting RUNX2. The study revealed 34 pathways regulated by miRNA-10 that directly affect RUNX2 expression. The extensive network associated with miR-10, in contrast to miR-9, indicates that miR-10 functions as a superior integrator, orchestrating several signals to culminate in RUNX2 activation.
g. Figure 7. Regulation of miRNA-9 Expression Mediated by PDLIM3. Network analysis identifies 49 pathways directly regulated by PDLIM3 that influence the expression of miRNA-9. This establishes a molecular connection, demonstrating how the structural protein PDLIM3 operates as a signaling hub to regulate epigenetic processes through miR-9.
h. Figure 8. PDLIM3 as a Principal Upstream Regulator of miRNA-10 Expression. A significant network of 98 pathways demonstrates direct regulation of miRNA-10 expression by PDLIM3. This signifies the most comprehensive connectivity in our investigation, pinpointing the control of miR-10 as a primary downstream function of PDLIM3 in the estrogen signaling pathway.
i. Figure 9. Integrated Molecular Network of the Estrogen-PDLIM3-miRNA-RUNX2 Axis. A fundamental network comprising 39 molecules that are directly or indirectly implicated in RUNX2 expression, as modulated by the interaction between estrogen, PDLIM3, miRNA-9, and miRNA-10. This integrated map vividly delineates the novel multi-layered regulatory pathway from estrogen receptor stimulation to the transcriptional regulation of RUNX2.
j. Figure 10. Innovative Network Incorporating miR-6769b in Estrogen-Driven Osteogenesis. The analysis revealed a network of 45 molecules that regulate RUNX2 expression, influenced by estrogen, PDLIM3, miR-1896, and the recently identified microRNA-6769B. This substantially broadens the suggested regulatory framework by incorporating miR-6769b as a new pivotal component.

---

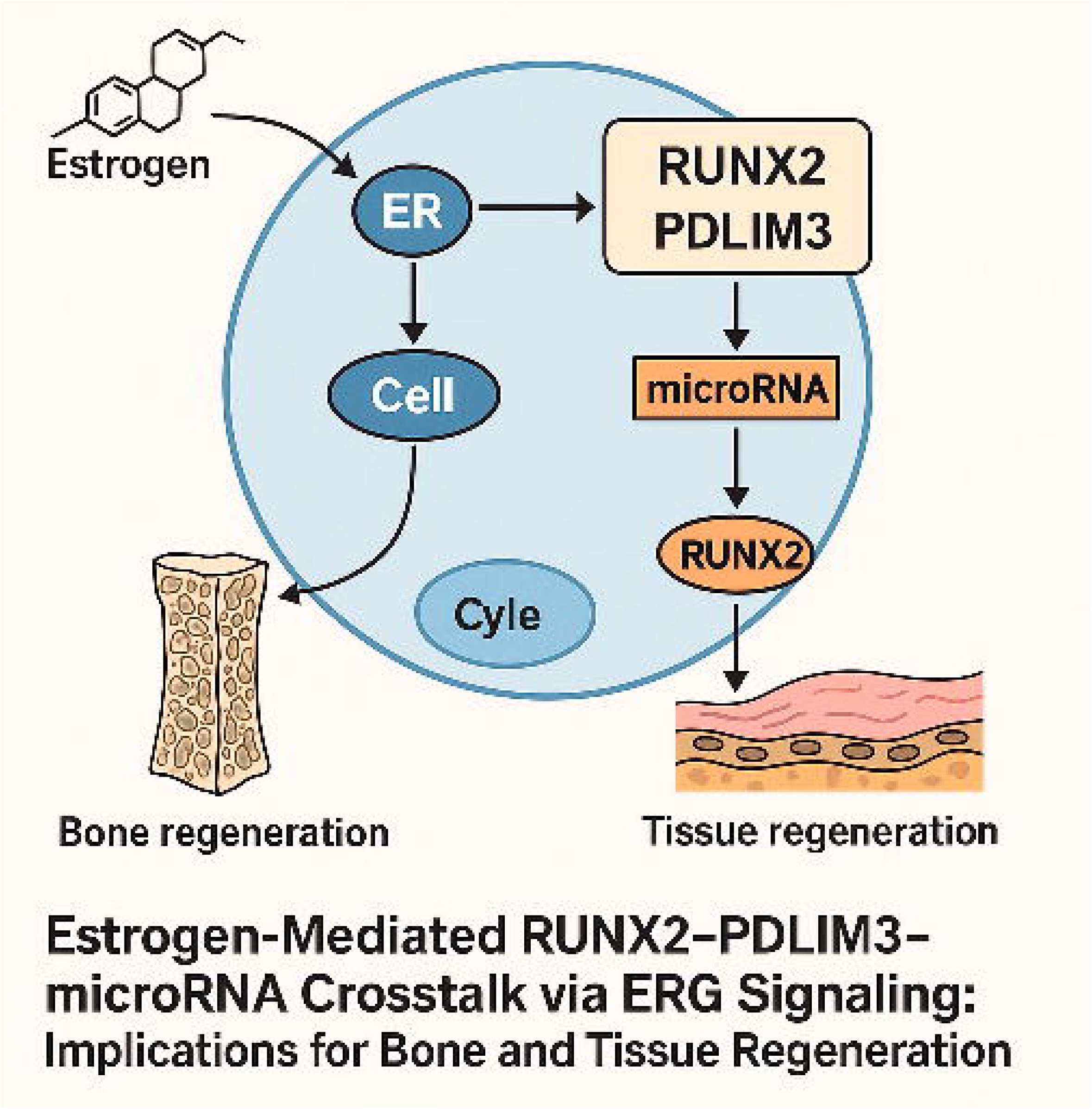

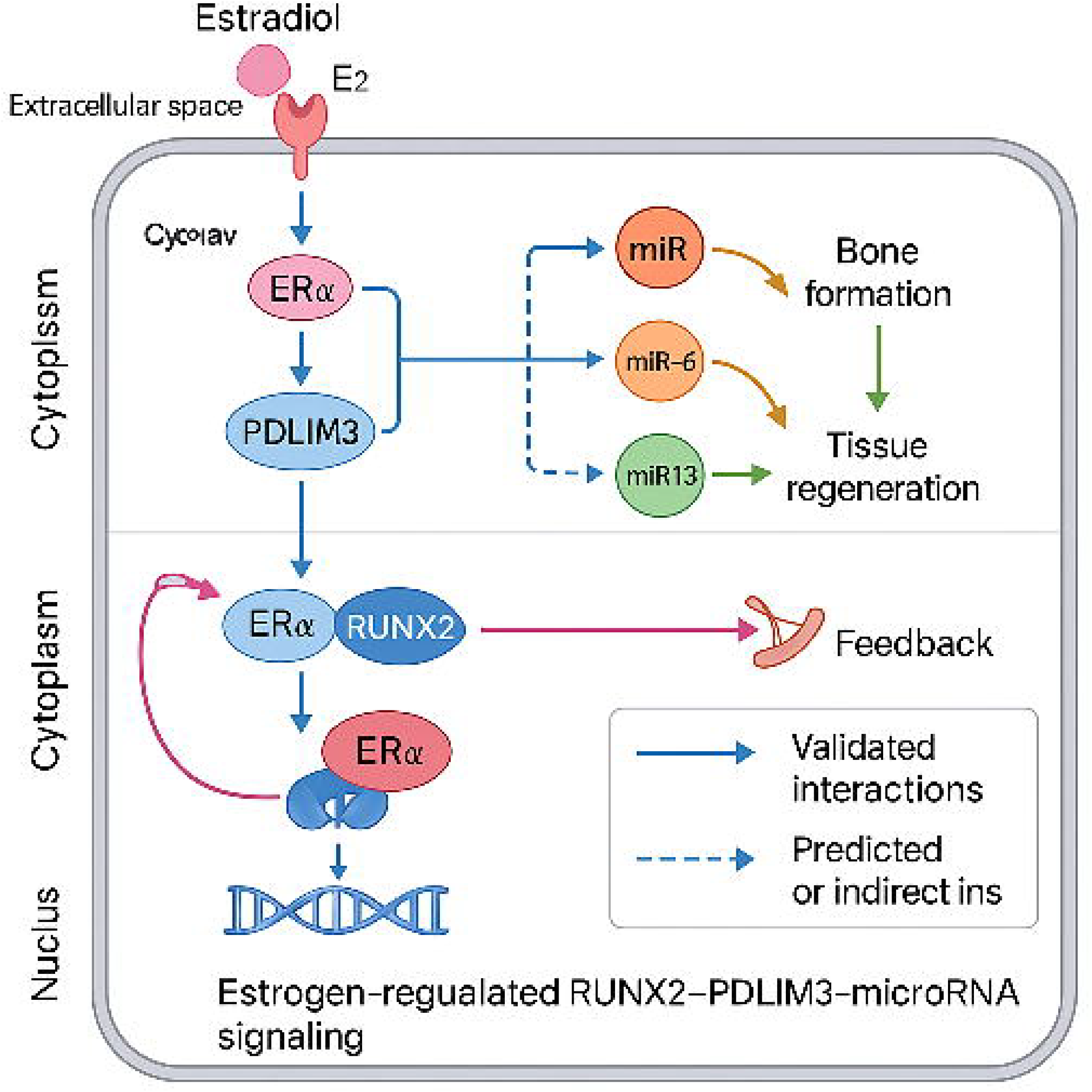

