## Supplementary material for "“Integrative Network Meta-Analysis Reveals Estrogen-Mediated RUNX2–PDLIM3–microRNA Crosstalk via ERG Signaling: Implications for Bone and Tissue Regeneration”": https://doi.org/10.5281/zenodo.17089993

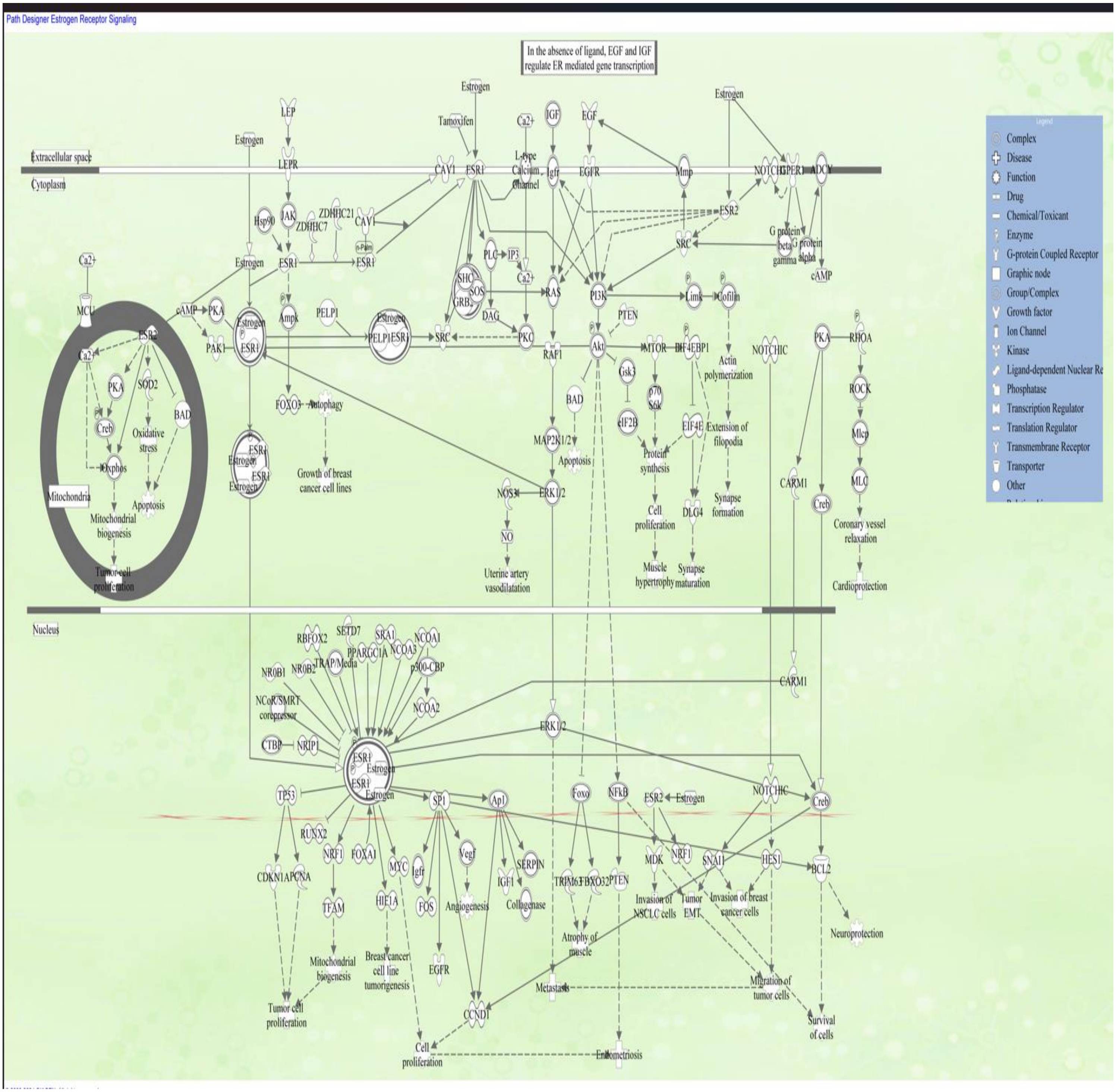


Figure1: Estrogen receptor signaling, designed and created using QIAGEN-IPA

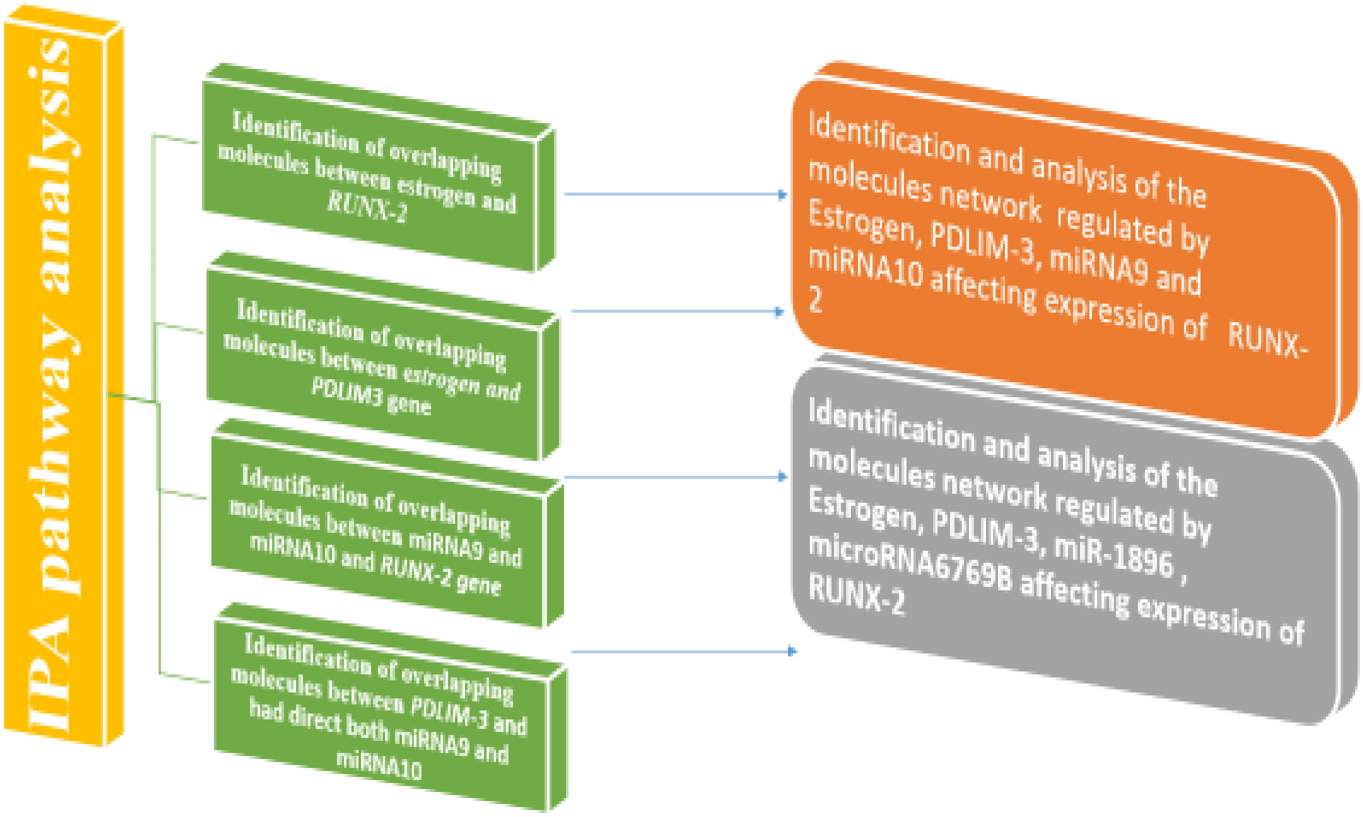


Figure.2: The data mining workflow was based on QIAGEN's Ingenuity Pathway Analysis (IPA) bioinformatics tools. The "Grow", "Connect", "Pathway Explorer", and "Molecule Activity Predictor" (MAP) tools from the "My Pathway" feature were used to develop biological networks that showed the connectivity between distinct nodes. Furthermore, the "Core Analysis: Expression Analysis" tool was utilized to compare the molecules within the produced molecular route to canonical pathways recorded within QIAGEN's knowledge base (QKB).

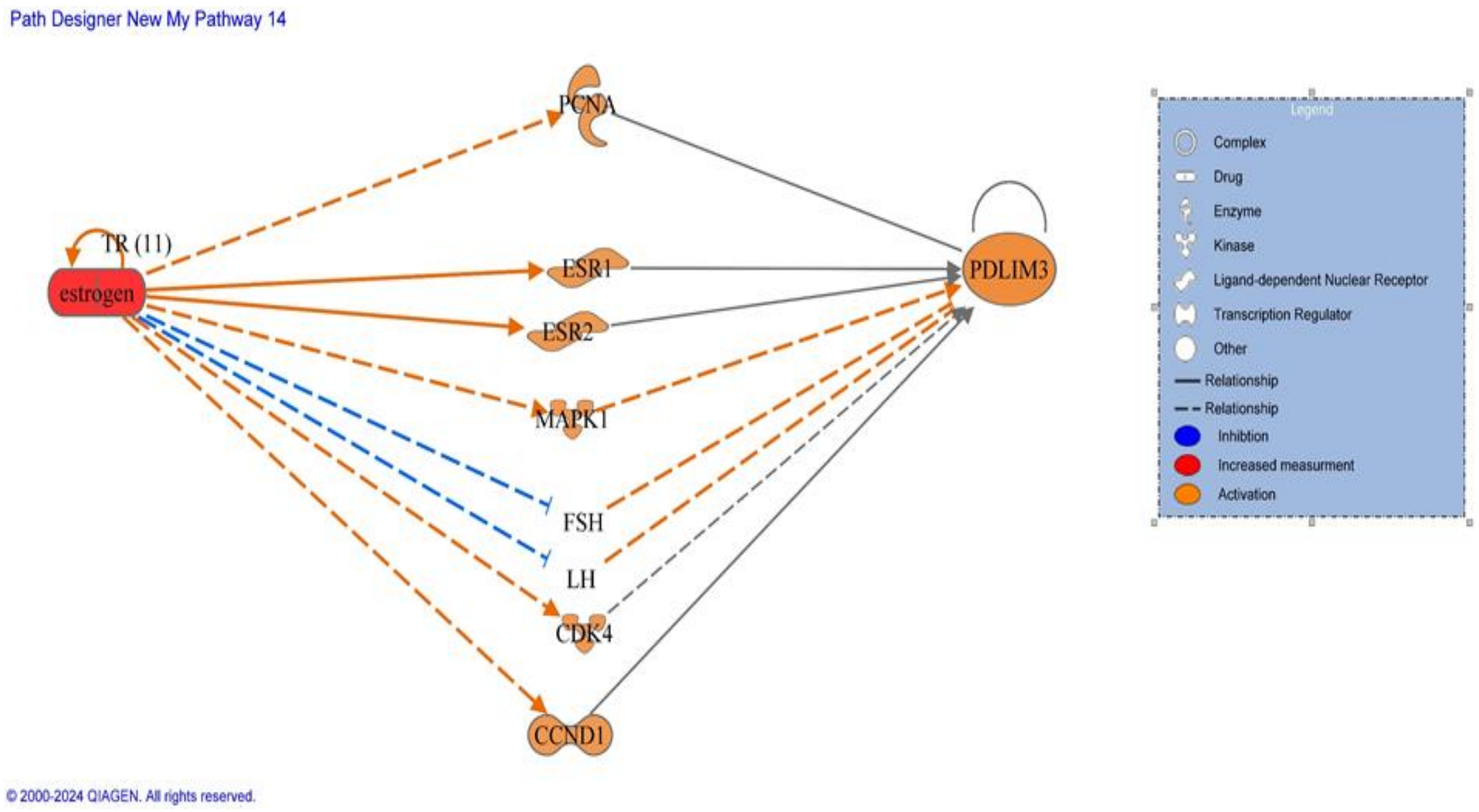


Figure.3: Molecular network depicting the connectivity and relationships among the overlapping molecules associated with direct influence of estrogen on PDLIM3 expression

Table.1 : Estrogen-regulated compounds that impact the expression of the *PDLIM3* gene

| Symbol | Gene Name | Location | Family |
|---|---|---|---|
| CCND1 | cyclin D1 | Nucleus | transcription regulator |
| CDK4 | cyclin dependent kinase 4 | Nucleus | kinase |
| ESR1 | estrogen receptor 1 | Nucleus | ligand-dependent nuclear receptor |
| ESR2 | estrogen receptor 2 | Nucleus | ligand-dependent nuclear receptor |
| Estrogen | estrogen | Other | chemical drug |
| FSH | Follicle stimulating hormone | Plasma Membrane | complex |
| LH | Luteliizng hormone | Plasma Membrane | Complex |
| MAPK1 | mitogen-activated protein kinase 1 | Cytoplasm | Kinase |
| PCNA | proliferating cell nuclear antigen | Nucleus | Enzyme |
| PDLIM3 | PDZ and LIM domain 3 | Cytoplasm | Other |

Table 2: Various interactions between molecules incorporated in the estrogen-*PDLIM3* pathway

| From Molecule(s) | Relationship Type | To Molecule(s) |
|---|---|---|
| 1. CCND1 | expression | PDLIM3 |
| 2. CDK4 | expression | PDLIM3 |
| 3. ESR1 | chemical-protein interactions | estrogen |
| 4. ESR1 | expression | PDLIM3 |
| 5. ESR1 | regulation of binding | Estrogen |
| 6. ESR2 | chemical-protein interactions | Estrogen |
| 7. ESR2 | expression | PDLIM3 |
| 8. FSH | expression | PDLIM3 |
| 9. LH | expression | PDLIM3 |
| 10. MAPK1 | expression | PDLIM3 |
| 11. PDLIM3 | protein-protein interactions | PCNA |
| 12. PDLIM3 | protein-protein interactions | PDLIM3 |
| 13. estrogen | activation | CDK4 |
| 14. estrogen | activation | ESR1 |
| 15. estrogen | activation | ESR2 |
| 16. estrogen | activation | MAPK1 |
| 17. estrogen | chemical-protein interactions | ESR1 |
| 18. estrogen | chemical-protein interactions | ESR2 |

| | | |
|---|---|---|
| 19. estrogen | expression | CCND1 |
| 20. estrogen | expression | ESR1 |
| 21. estrogen | expression | ESR2 |
| 22. estrogen | expression | FSH |
| 23. estrogen | expression | LH |
| 24. estrogen | expression | PCNA |
| 25. estrogen | localization | FSH |
| 26. estrogen | localization | LH |
| 27. estrogen | molecular cleavage | ESR1 |
| 28. estrogen | phosphorylation | ESR1 |
| 29. estrogen | phosphorylation | ESR2 |
| 30. estrogen | phosphorylation | MAPK1 |
| 31. estrogen | regulation of binding | CCND1 |
| 32. estrogen | regulation of binding | CDK4 |
| 33. estrogen | regulation of binding | ESR1 |
| 34. estrogen | regulation of binding | ESR2 |
| 35. estrogen | transcription | CCND1 |
| 36. estrogen | translocation | ESR1 |
| 37. estrogen | translocation | ESR2 |
| 38. estrogen | translocation | estrogen |

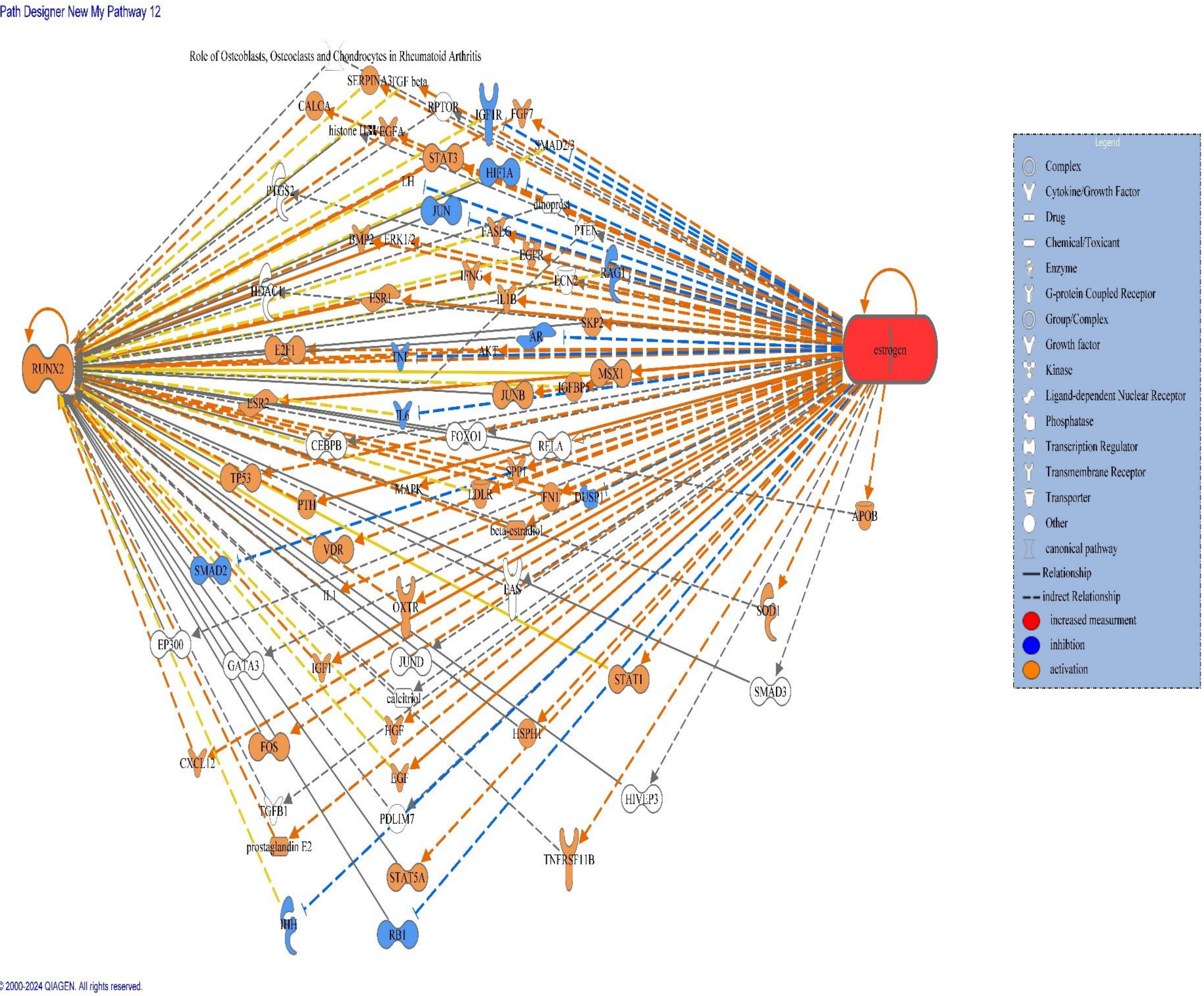


Figure.4: Molecular network depicting the connectivity and relationships among the overlapping molecules associated with direct influence of estrogen on *RUNX-2* expression

Table.3: Estrogen-regulated compounds that impact the expression of the *RUNX-2* gene

| Symbol | molecule/ Gene Name | Location | Family |
|---|---|---|---|
| 39. AKT | AKT Serine/Threonine Kinase 1. | Cytoplasm | group |
| 40. APOB | apolipoprotein B | Extracellular Space | Transporter |
| 41. AR | androgen receptor | Nucleus | ligand-dependent nuclear receptor |
| 42. beta-estradiol | beta-estradiol | Other | chemical - endogenous mammalian |
| 43. BMP2 | bone morphogenetic protein 2 | Extracellular Space | growth factor |
| 44. CALCA | calcitonin related polypeptide alpha | Plasma Membrane | Other |
| 45. Calcitriol | calcitriol | Other | chemical drug |
| 46. CEBPB | CCAAT enhancer binding protein beta | Nucleus | transcription regulator |
| 47. CXCL12 | C-X-C motif chemokine ligand 12 | Extracellular Space | Cytokine |
| 48. dinoprost | dinoprost | Other | chemical - endogenous mammalian |
| 49. DUSP1 | dual specificity phosphatase 1 | Nucleus | Phosphatase |
| 50. E2F1 | E2F transcription factor 1 | Nucleus | transcription regulator |
| 51. EGF | epidermal growth factor | Extracellular Space | growth factor |
| 52. EGFR | epidermal growth factor receptor | Plasma Membrane | Kinase |
| 53. EP300 | E1A binding protein p300 | Nucleus | transcription regulator |
| 54. ERK1/2 | ERK1/2 | Cytoplasm | Group |
| 55. ESR1 | estrogen receptor 1 | Nucleus | ligand-dependent nuclear receptor |
| 56. ESR2 | estrogen receptor 2 | Nucleus | ligand-dependent nuclear receptor |
| 57. Estrogen | estrogen | Other | chemical drug |
| 58. FAS | Fas cell surface death receptor | Plasma Membrane | transmembrane receptor |
| 59. FASLG | Fas ligand | Extracellular Space | Cytokine |
| 60. FGF7 | fibroblast growth factor 7 | Extracellular Space | growth factor |
| 61. FN1 | fibronectin 1 | Extracellular Space | Other |
| 62. FOS | Fos proto-oncogene, AP-1 transcription factor subunit | Nucleus | transcription regulator |
| 63. FOXO1 | forkhead box O1 | Nucleus | transcription regulator |

| | | | |
|---|---|---|---|
| 64. GATA3 | GATA binding protein 3 | Nucleus | transcription regulator |
| 65. HDAC1 | histone deacetylase 1 | Nucleus | enzyme |
| 66. HGF | hepatocyte growth factor | Extracellular Space | growth factor |
| 67. HIF1A | hypoxia inducible factor 1 subunit alpha | Nucleus | transcription regulator |
| 68. histone H3 | histone H3 | Nucleus | group |
| 69. HIVEP3 | HIVEP zinc finger 3 | Nucleus | transcription regulator |
| 70. HSPH1 | heat shock protein family H (Hsp110) member 1 | Cytoplasm | other |
| 71. IFNG | interferon gamma | Extracellular Space | cytokine |
| 72. IGF1 | insulin like growth factor 1 | Extracellular Space | growth factor |
| 73. IGF1R | insulin like growth factor 1 receptor | Plasma Membrane | transmembrane receptor |
| 74. IGFBP5 | insulin like growth factor binding protein 5 | Extracellular Space | other |
| 75. IHH | Indian hedgehog signaling molecule | Extracellular Space | enzyme |
| 76. IL1 | IL1 | Extracellular Space | group |
| 77. IL1B | interleukin 1 beta | Extracellular Space | cytokine |
| 78. IL6 | interleukin 6 | Extracellular Space | cytokine |
| 79. JUN | Jun proto-oncogene, AP-1 transcription factor subunit | Nucleus | transcription regulator |
| 80. JUNB | JunB proto-oncogene, AP-1 transcription factor subunit | Nucleus | transcription regulator |
| 81. JUND | JunD proto-oncogene, AP-1 transcription factor subunit | Nucleus | transcription regulator |
| 82. LCN2 | lipocalin 2 | Extracellular Space | transporter |
| 83. LDLR | low density lipoprotein receptor | Plasma Membrane | Transporter |
| 84. LH | LH | Plasma Membrane | Complex |
| 85. MAPK | MAPK | Cytoplasm | Group |
| 86. MSX1 | msh homeobox 1 | Nucleus | transcription regulator |
| 87. OXTR | oxytocin receptor | Plasma Membrane | G-protein coupled receptor |
| 88. PDLIM7 | PDZ and LIM domain 7 | Cytoplasm | Other |
| 89. prostaglandin E2 | prostaglandin E2 | Other | chemical - endogenous mammalian |
| 90. PTEN | phosphatase and tensin homolog | Cytoplasm | Phosphatase |
| 91. PTGS2 | prostaglandin-endoperoxide synthase 2 | Cytoplasm | Enzyme |
| 92. PTH | parathyroid hormone | Extracellular Space | Other |

| | | | |
|---|---|---|---|
| 93. RAG1 | recombination activating 1 | Nucleus | Enzyme |
| 94. RB1 | RB transcriptional corepressor 1 | Nucleus | transcription regulator |
| 95. RELA | RELA proto-oncogene, NF-kB subunit | Nucleus | transcription regulator |
| 96. RPTOR | regulatory associated protein of MTOR complex 1 | Cytoplasm | Other |
| 97. RUNX2 | RUNX family transcription factor 2 | Nucleus | transcription regulator |
| 98. SERPINA3 | serpin family A member 3 | Extracellular Space | Other |
| 99. SKP2 | S-phase kinase associated protein 2 | Nucleus | other |
| 100. SMAD2 | SMAD family member 2 | Nucleus | transcription regulator |
| 101. SMAD2/3 | SMAD2/3 | Cytoplasm | group |
| 102. SMAD3 | SMAD family member 3 | Nucleus | transcription regulator |
| 103. SOD1 | superoxide dismutase 1 | Cytoplasm | Enzyme |
| 104. SPP1 | secreted phosphoprotein 1 | Extracellular Space | Cytokine |
| 105. STAT1 | signal transducer and activator of transcription 1 | Nucleus | transcription regulator |
| 106. STAT3 | signal transducer and activator of transcription 3 | Nucleus | transcription regulator |
| 107. STAT5A | signal transducer and activator of transcription 5A | Nucleus | transcription regulator |
| 108. TGF beta | TGF beta | Extracellular Space | Group |
| 109. TGFB1 | transforming growth factor beta 1 | Extracellular Space | growth factor |
| 110. TNF | tumor necrosis factor | Extracellular Space | Cytokine |
| 111. TNFRSF11B | TNF receptor superfamily member 11b | Plasma Membrane | transmembrane receptor |
| 112. TP53 | tumor protein p53 | Nucleus | transcription regulator |
| 113. VDR | vitamin D receptor | Nucleus | transcription regulator |
| 114. VEGFA | vascular endothelial growth factor A | Extracellular Space | growth factor |

Table4: Various interactions between molecules incorporated in the Estrogen-*RUNX-2* pathway

| From Molecule(s) | Relationship Type | To Molecule(s) |
|---|---|---|
| 1. AR | protein-protein interactions | ESR2 |
| 2. AR | protein-protein interactions | FKBP4 |
| 3. AR | protein-protein interactions | HSP90 (family) |
| 4. ASAH1 | expression | RUNX2 |
| 5. CBL | activation | RUNX2 |
| 6. CBL | expression | RUNX2 |
| 7. CBX5 | expression | RUNX2 |
| 8. CEBPA | expression | RUNX2 |
| 9. CEBPB | expression | RUNX2 |
| 10. CEBPB | protein-protein interactions | ESR1 |
| 11. CEBPB | protein-protein interactions | RUNX2 |
| 12. CREBBP | expression | RUNX2 |
| 13. CREBBP | protein-protein interactions | AFP |
| 14. CREBBP | protein-protein interactions | RUNX2 |
| 15. CTNNB1 | expression | RUNX2 |
| 16. CTNNB1 | inhibition | RUNX2 |
| 17. CTNNB1 | protein-DNA interactions | RUNX2 |
| 18. CTNNB1 | protein-protein interactions | RUNX2 |
| 19. CTNNB1 | transcription | RUNX2 |
| 20. DDX5 | protein-protein interactions | ESR1 |
| 21. EED | expression | RUNX2 |
| 22. EED | modification | RUNX2 |
| 23. ELK1 | expression | RUNX2 |
| 24. ELK1 | protein-DNA interactions | RUNX2 |
| 25. EP300 | expression | RUNX2 |
| 26. EP300 | protein-protein interactions | ESR1 |
| 27. EP300 | protein-protein interactions | RUNX2 |
| 28. ER/estrogen | membership | ESR1 |
| 29. ERBB2 | activation | CTNNB1 |
| 30. ERBB2 | activation | EP300 |
| 31. ERBB2 | activation | ERK1/2 |
| 32. ERBB2 | activation | NOTCH1 |
| 33. ERBB2 | activation | SP1 |
| 34. ERBB2 | activation | STAT1 |
| 35. ERBB2 | activation | STAT3 |
| 36. ERBB2 | activation | histone H3 |
| 37. ERBB2 | inhibition | FOXO3 |
| 38. ERBB2 | inhibition | RB |
| 39. ERBB2 | inhibition | RB1 |
| 40. ERBB2 | phosphorylation | CTNNB1 |

| | | |
|---|---|---|
| 41. ERBB2 | phosphorylation | EP300 |
| 42. ERBB2 | phosphorylation | ERK1/2 |
| 43. ERBB2 | phosphorylation | FOXO3 |
| 44. ERBB2 | phosphorylation | RB |
| 45. ERBB2 | phosphorylation | RB1 |
| 46. ERBB2 | phosphorylation | SP1 |
| 47. ERBB2 | phosphorylation | STAT1 |
| 48. ERBB2 | phosphorylation | STAT3 |
| 49. ERBB2 | phosphorylation | histone H3 |
| 50. ERBB2 | protein-protein interactions | CBL |
| 51. ERBB2 | protein-protein interactions | CCNB1 |
| 52. ERBB2 | protein-protein interactions | CTNNB1 |
| 53. ERBB2 | protein-protein interactions | CUL4A |
| 54. ERBB2 | protein-protein interactions | NOTCH1 |
| 55. ERBB2 | protein-protein interactions | STAT1 |
| 56. ERBB2 | protein-protein interactions | STAT3 |
| 57. ERK1/2 | activation | RUNX2 |
| 58. ERK1/2 | protein-protein interactions | RUNX2 |
| 59. ESR1 | RNA-RNA interactions: non-targeting interactions | ESR2 |
| 60. ESR1 | activation | Ap1 |
| 61. ESR1 | activation | ESR2 |
| 62. ESR1 | activation | RELA |
| 63. ESR1 | activation | RUNX2 |
| 64. ESR1 | activation | SMAD3 |
| 65. ESR1 | activation | SP1 |
| 66. ESR1 | activation | STAT5A |
| 67. ESR1 | activation | TP53 |
| 68. ESR1 | chemical-protein interactions | estrogen |
| 69. ESR1 | expression | AR |
| 70. ESR1 | expression | CBL |
| 71. ESR1 | expression | CCNB1 |
| 72. ESR1 | expression | CEBPA |
| 73. ESR1 | expression | CEBPB |
| 74. ESR1 | expression | CEBPD |
| 75. ESR1 | expression | CTNNB1 |
| 76. ESR1 | expression | CUL4B |
| 77. ESR1 | expression | EHMT2 |
| 78. ESR1 | expression | EP300 |
| 79. ESR1 | expression | ESR2 |
| 80. ESR1 | expression | FOS |
| 81. ESR1 | expression | FOSB |
| 82. ESR1 | expression | FOSL1 |
| 83. ESR1 | expression | FOSL2 |

| | | |
|---|---|---|
| 84. ESR1 | expression | FOXO1 |
| 85. ESR1 | expression | FOXO4 |
| 86. ESR1 | expression | GATA3 |
| 87. ESR1 | expression | GLI2 |
| 88. ESR1 | expression | GSN |
| 89. ESR1 | expression | HES1 |
| 90. ESR1 | expression | HEY1 |
| 91. ESR1 | expression | HIF1A |
| 92. ESR1 | expression | HSPD1 |
| 93. ESR1 | expression | HSPH1 |
| 94. ESR1 | expression | ID1 |
| 95. ESR1 | expression | IRF4 |
| 96. ESR1 | expression | JUN |
| 97. ESR1 | expression | JUNB |
| 98. ESR1 | expression | JUND |
| 99. ESR1 | expression | LIMA1 |
| 100. ESR1 | expression | NFYB |
| 101. ESR1 | expression | NOTCH1 |
| 102. ESR1 | expression | NR0B2 |
| 103. ESR1 | expression | OSTF1 |
| 104. ESR1 | expression | PPARD |
| 105. ESR1 | expression | PPARG |
| 106. ESR1 | expression | RB1 |
| 107. ESR1 | expression | RBL2 |
| 108. ESR1 | expression | RELA |
| 109. ESR1 | expression | RUNX2 |
| 110. ESR1 | expression | SKP2 |
| 111. ESR1 | expression | SMAD2 |
| 112. ESR1 | expression | SMAD3 |
| 113. ESR1 | expression | SMAD5 |
| 114. ESR1 | expression | SMAD6 |
| 115. ESR1 | expression | SMURF1 |
| 116. ESR1 | expression | SMURF2 |
| 117. ESR1 | expression | SNAI1 |
| 118. ESR1 | expression | SNAI2 |
| 119. ESR1 | expression | SOX9 |
| 120. ESR1 | expression | SP1 |
| 121. ESR1 | expression | STAT1 |
| 122. ESR1 | expression | STAT3 |
| 123. ESR1 | expression | STAT5A |
| 124. ESR1 | expression | TCF7L2 |
| 125. ESR1 | expression | THRB |
| 126. ESR1 | expression | TP53 |
| 127. ESR1 | expression | TRIB3 |

| | | | |
|---|---|---|---|
| 128. | ESR1 | expression | TSC22D3 |
| 129. | ESR1 | expression | VDR |
| 130. | ESR1 | expression | ZBTB7B |
| 131. | ESR1 | inhibition | RELA |
| 132. | ESR1 | inhibition | RUNX2 |
| 133. | ESR1 | inhibition | TP53 |
| 134. | ESR1 | protein-DNA interactions | CREBBP |
| 135. | ESR1 | protein-DNA interactions | ESRRA |
| 136. | ESR1 | protein-DNA interactions | FOSL1 |
| 137. | ESR1 | protein-DNA interactions | FOXC1 |
| 138. | ESR1 | protein-DNA interactions | GATA3 |
| 139. | ESR1 | protein-DNA interactions | KAT6B |
| 140. | ESR1 | protein-DNA interactions | NR0B2 |
| 141. | ESR1 | protein-DNA interactions | PPARGC1A |
| 142. | ESR1 | protein-DNA interactions | SIRT1 |
| 143. | ESR1 | protein-DNA interactions | SOX9 |
| 144. | ESR1 | protein-DNA interactions | STAT5A |
| 145. | ESR1 | protein-DNA interactions | TP53 |
| 146. | ESR1 | protein-DNA interactions | ZMYND8 |
| 147. | ESR1 | protein-protein interactions | ALYREF |
| 148. | ESR1 | protein-protein interactions | AR |
| 149. | ESR1 | protein-protein interactions | Ap1 |
| 150. | ESR1 | protein-protein interactions | CBX5 |
| 151. | ESR1 | protein-protein interactions | CEBPA |
| 152. | ESR1 | protein-protein interactions | CEBPB |
| 153. | ESR1 | protein-protein interactions | CIC |
| 154. | ESR1 | protein-protein interactions | CREBBP |
| 155. | ESR1 | protein-protein interactions | CTBP2 |
| 156. | ESR1 | protein-protein interactions | CTNNB1 |
| 157. | ESR1 | protein-protein interactions | CUL4B |
| 158. | ESR1 | protein-protein interactions | DDX5 |
| 159. | ESR1 | protein-protein interactions | EHMT2 |
| 160. | ESR1 | protein-protein interactions | EP300 |
| 161. | ESR1 | protein-protein interactions | ERBB2 |
| 162. | ESR1 | protein-protein interactions | ESR2 |
| 163. | ESR1 | protein-protein interactions | ESRRA |
| 164. | ESR1 | protein-protein interactions | FOS |
| 165. | ESR1 | protein-protein interactions | FOSL2 |
| 166. | ESR1 | protein-protein interactions | FOXO1 |
| 167. | ESR1 | protein-protein interactions | FOXO4 |
| 168. | ESR1 | protein-protein interactions | GATA3 |
| 169. | ESR1 | protein-protein interactions | GSN |
| 170. | ESR1 | protein-protein interactions | HDAC1 |
| 171. | ESR1 | protein-protein interactions | HIF1A |

| | | | |
|---|---|---|---|
| 172. | ESR1 | protein-protein interactions | HSPD1 |
| 173. | ESR1 | protein-protein interactions | HSPH1 |
| 174. | ESR1 | protein-protein interactions | JUN |
| 175. | ESR1 | protein-protein interactions | JUNB |
| 176. | ESR1 | protein-protein interactions | JUND |
| 177. | ESR1 | protein-protein interactions | LIMA1 |
| 178. | ESR1 | protein-protein interactions | NR0B2 |
| 179. | ESR1 | protein-protein interactions | PPARG |
| 180. | ESR1 | protein-protein interactions | PPARGC1A |
| 181. | ESR1 | protein-protein interactions | RELA |
| 182. | ESR1 | protein-protein interactions | RUNX2 |
| 183. | ESR1 | protein-protein interactions | SIRT1 |
| 184. | ESR1 | protein-protein interactions | SKP2 |
| 185. | ESR1 | protein-protein interactions | SMAD2 |
| 186. | ESR1 | protein-protein interactions | SMAD3 |
| 187. | ESR1 | protein-protein interactions | SMURF1 |
| 188. | ESR1 | protein-protein interactions | SP1 |
| 189. | ESR1 | protein-protein interactions | STAT1 |
| 190. | ESR1 | protein-protein interactions | STAT3 |
| 191. | ESR1 | protein-protein interactions | STAT5A |
| 192. | ESR1 | protein-protein interactions | TCF7L2 |
| 193. | ESR1 | protein-protein interactions | TP53 |
| 194. | ESR1 | protein-protein interactions | ZBTB7B |
| 195. | ESR1 | protein-protein interactions | ZMYND8 |
| 196. | ESR1 | transcription | ESRRA |
| 197. | ESR1 | transcription | FOS |
| 198. | ESR1 | transcription | PPARGC1A |
| 199. | ESR1 | transcription | RUNX2 |
| 200. | ESR1 | transcription | SIRT1 |
| 201. | ESR1 | transcription | TP53 |
| 202. | ESR2 | RNA-RNA interactions: non-targeting interactions | ESR1 |
| 203. | ESR2 | activation | AR |
| 204. | ESR2 | activation | ESR1 |
| 205. | ESR2 | activation | RELA |
| 206. | ESR2 | activation | SP1 |
| 207. | ESR2 | activation | STAT3 |
| 208. | ESR2 | activation | STAT5A |
| 209. | ESR2 | chemical-protein interactions | estrogen |
| 210. | ESR2 | expression | AR |
| 211. | ESR2 | expression | CEBPD |
| 212. | ESR2 | expression | CREBBP |
| 213. | ESR2 | expression | CTNNB1 |
| 214. | ESR2 | expression | EHMT2 |

| | | | |
|---|---|---|---|
| 215. | ESR2 | expression | ELK1 |
| 216. | ESR2 | expression | EP300 |
| 217. | ESR2 | expression | ESR1 |
| 218. | ESR2 | expression | EZH2 |
| 219. | ESR2 | expression | FOS |
| 220. | ESR2 | expression | FOSL2 |
| 221. | ESR2 | expression | FOXC2 |
| 222. | ESR2 | expression | FOXO1 |
| 223. | ESR2 | expression | FOXO3 |
| 224. | ESR2 | expression | GATA1 |
| 225. | ESR2 | expression | GATA3 |
| 226. | ESR2 | expression | HSPD1 |
| 227. | ESR2 | expression | JAG2 |
| 228. | ESR2 | expression | JUNB |
| 229. | ESR2 | expression | KLF4 |
| 230. | ESR2 | expression | NOTCH1 |
| 231. | ESR2 | expression | RELA |
| 232. | ESR2 | expression | RUNX2 |
| 233. | ESR2 | expression | SKP2 |
| 234. | ESR2 | expression | SMAD2 |
| 235. | ESR2 | expression | SMAD3 |
| 236. | ESR2 | expression | SMAD4 |
| 237. | ESR2 | expression | SNAI1 |
| 238. | ESR2 | expression | SNAI2 |
| 239. | ESR2 | expression | SP1 |
| 240. | ESR2 | expression | SRF |
| 241. | ESR2 | expression | TWIST1 |
| 242. | ESR2 | expression | YY1 |
| 243. | ESR2 | expression | osteocalcin |
| 244. | ESR2 | inhibition | RELA |
| 245. | ESR2 | protein-DNA interactions | FOS |
| 246. | ESR2 | protein-RNA interactions | CTNNB1 |
| 247. | ESR2 | protein-protein interactions | ALYREF |
| 248. | ESR2 | protein-protein interactions | AR |
| 249. | ESR2 | protein-protein interactions | ASAH1 |
| 250. | ESR2 | protein-protein interactions | CIC |
| 251. | ESR2 | protein-protein interactions | CREBBP |
| 252. | ESR2 | protein-protein interactions | CTBP2 |
| 253. | ESR2 | protein-protein interactions | CTNNB1 |
| 254. | ESR2 | protein-protein interactions | EED |
| 255. | ESR2 | protein-protein interactions | EHMT2 |
| 256. | ESR2 | protein-protein interactions | EP300 |
| 257. | ESR2 | protein-protein interactions | ESR1 |
| 258. | ESR2 | protein-protein interactions | FOS |

| | | |
|---|---|---|
| 259. ESR2 | protein-protein interactions | FOXO3 |
| 260. ESR2 | protein-protein interactions | HSPD1 |
| 261. ESR2 | protein-protein interactions | SMAD3 |
| 262. ESR2 | protein-protein interactions | SMAD4 |
| 263. ESR2 | protein-protein interactions | SP1 |
| 264. ESR2 | protein-protein interactions | STAT3 |
| 265. ESR2 | protein-protein interactions | STAT5A |
| 266. ESRRA | protein-DNA interactions | RUNX2 |
| 267. ESRRA | protein-protein interactions | ESR1 |
| 268. ESRRA | transcription | RUNX2 |
| 269. ESTG:ESR1:chaperone | membership | ESR1 |
| 270. ESTG:ESR2:chaperone | membership | ESR2 |
| 271. ESTG:Me-PalmS-ESR dimers | membership | ESR1 |
| 272. ESTG:Me-PalmS-ESR dimers | membership | ESR2 |
| 273. ETS1 | protein-DNA interactions | RUNX2 |
| 274. ETS1 | protein-protein interactions | ESR1 |
| 275. ETS1 | protein-protein interactions | RUNX2 |
| 276. ETS1 | transcription | RUNX2 |
| 277. EWSR1 | protein-protein interactions | ESR1 |
| 278. EZH2 | expression | RUNX2 |
| 279. EZH2 | protein-protein interactions | ESR1 |
| 280. FHL2 | protein-protein interactions | ESR1 |
| 281. FHL2 | protein-protein interactions | ESR2 |
| 282. FKBP4 | activation | AR |
| 283. FKBP4 | chemical-protein interactions | estrogen |
| 284. FKBP4 | protein-protein interactions | AR |
| 285. FKBP4 | protein-protein interactions | CTNNB1 |
| 286. FKBP4 | protein-protein interactions | DET1 |
| 287. FKBP4 | protein-protein interactions | ESR1 |
| 288. FKBP4 | protein-protein interactions | EWSR1 |
| 289. FKBP4 | protein-protein interactions | EZH2 |
| 290. FKBP4 | protein-protein interactions | NR3C1 |
| 291. FOS | protein-DNA interactions | RUNX2 |
| 292. FOS | protein-protein interactions | ESR1 |
| 293. FOS | protein-protein interactions | ESR2 |
| 294. FOS | protein-protein interactions | RUNX2 |
| 295. FOS | protein-protein interactions | estrogen receptor |
| 296. FOSB | protein-DNA interactions | RUNX2 |
| 297. FOSL1 | protein-DNA interactions | RUNX2 |
| 298. FOSL2 | protein-DNA interactions | RUNX2 |
| 299. FOSL2 | protein-protein interactions | ESR1 |
| 300. FOXC1 | expression | RUNX2 |

| | | |
|---|---|---|
| 301. FOXC2 | expression | RUNX2 |
| 302. FOXO1 | activation | RUNX2 |
| 303. FOXO1 | expression | RUNX2 |
| 304. FOXO1 | protein-protein interactions | ESR1 |
| 305. FOXO1 | protein-protein interactions | RUNX2 |
| 306. FOXO1 | protein-protein interactions | estrogen receptor |
| 307. FOXO3 | expression | RUNX2 |
| 308. FOXO3 | protein-protein interactions | ESR1 |
| 309. FOXO3 | protein-protein interactions | ESR2 |
| 310. FOXO3 | protein-protein interactions | FKBP4 |
| 311. FOXO4 | expression | RUNX2 |
| 312. FOXO4 | protein-protein interactions | ESR1 |
| 313. GATA1 | expression | RUNX2 |
| 314. GATA3 | protein-DNA interactions | ESR1 |
| 315. GATA3 | protein-protein interactions | ESR1 |
| 316. GLI2 | protein-protein interactions | RUNX2 |
| 317. GLI2 | transcription | RUNX2 |
| 318. GNB2 | protein-protein interactions | ESR1 |
| 319. GNB2 | protein-protein interactions | ESR2 |
| 320. GNB2 | protein-protein interactions | FKBP4 |
| 321. GPER1:Heterotrimeric G-protein Gs:ESTG | membership | GNB2 |
| 322. GSN | protein-protein interactions | ERBB2 |
| 323. GSN | protein-protein interactions | ESR1 |
| 324. GSN | protein-protein interactions | ESR2 |
| 325. HDAC1 | protein-protein interactions | ESR1 |
| 326. HDAC1 | protein-protein interactions | ESR2 |
| 327. HDAC3 | activation | RUNX2 |
| 328. HDAC3 | protein-protein interactions | ESR1 |
| 329. HDAC3 | protein-protein interactions | ESR2 |
| 330. HDAC3 | protein-protein interactions | RUNX2 |
| 331. HDAC4 | activation | RUNX2 |
| 332. HDAC4 | expression | RUNX2 |
| 333. HDAC4 | protein-protein interactions | ESR1 |
| 334. HDAC4 | protein-protein interactions | RUNX2 |
| 335. HDAC5 | expression | RUNX2 |
| 336. HDAC5 | protein-protein interactions | ESR1 |
| 337. HDAC5 | protein-protein interactions | RUNX2 |
| 338. HDAC6 | protein-protein interactions | ERBB2 |
| 339. HDAC6 | protein-protein interactions | ESR2 |
| 340. HDAC7 | protein-protein interactions | ESR1 |
| 341. HES1 | activation | RUNX2 |
| 342. HES1 | protein-protein interactions | RUNX2 |
| 343. HIF1A | protein-protein interactions | ESR1 |

| | | | |
|---|---|---|---|
| 344. | HMGB2 | expression | RUNX2 |
| 345. | HMGB2 | protein-DNA interactions | RUNX2 |
| 346. | HMGB2 | protein-protein interactions | ESR1 |
| 347. | HMGB2 | protein-protein interactions | RUNX2 |
| 348. | HMGB2 | protein-protein interactions | estrogen receptor |
| 349. | HMGB2 | transcription | RUNX2 |
| 350. | HSP90 (family) | activation | AR |
| 351. | HSP90 (family) | activation | STAT3 |
| 352. | HSP90 (family) | activation | TP53 |
| 353. | HSP90 (family) | chemical-protein interactions | estrogen |
| 354. | HSP90 (family) | protein-protein interactions | AR |
| 355. | HSP90 (family) | protein-protein interactions | ESR1 |
| 356. | HSP90 (family) | protein-protein interactions | EWSR1 |
| 357. | HSP90 (family) | protein-protein interactions | GATA3 |
| 358. | HSP90 (family) | protein-protein interactions | HDAC1 |
| 359. | HSP90 (family) | protein-protein interactions | HDAC6 |
| 360. | HSP90 (family) | protein-protein interactions | HIF1A |
| 361. | HSP90 (family) | protein-protein interactions | NR3C1 |
| 362. | HSP90 (family) | protein-protein interactions | STAT3 |
| 363. | HSP90 (family) | protein-protein interactions | STUB1 |
| 364. | HSP90 (family) | protein-protein interactions | TP53 |
| 365. | HSP90 (family) | protein-protein interactions | histone H4 |
| 366. | HSPA4 | protein-protein interactions | ERBB2 |
| 367. | HSPA4 | protein-protein interactions | ESR1 |
| 368. | HSPA4 | protein-protein interactions | ESR2 |
| 369. | HSPA4 | protein-protein interactions | FKBP4 |
| 370. | HSPA4L | protein-protein interactions | ERBB2 |
| 371. | HSPA4L | protein-protein interactions | ESR1 |
| 372. | HSPD1 | protein-protein interactions | ERBB2 |
| 373. | HSPD1 | protein-protein interactions | ESR1 |
| 374. | HSPD1 | protein-protein interactions | ESR2 |
| 375. | HSPH1 | protein-protein interactions | ESR1 |
| 376. | HSPH1 | protein-protein interactions | ESR2 |
| 377. | HSPH1 | protein-protein interactions | HSP90 (family) |
| 378. | ID1 | expression | RUNX2 |
| 379. | IFI16 | activation | RUNX2 |
| 380. | IFI16 | protein-protein interactions | FKBP4 |
| 381. | IFI16 | protein-protein interactions | RUNX2 |
| 382. | IGF1 | activation | ESR1 |
| 383. | IGF1 | activation | TP53 |
| 384. | IGF1 | chemical-protein interactions | estrogen |
| 385. | IMPDH1 | protein-protein interactions | ESR1 |
| 386. | IRF4 | expression | RUNX2 |
| 387. | IRF4 | protein-protein interactions | FKBP4 |

| | | | |
|---|---|---|---|
| 388. | JUN | protein-DNA interactions | RUNX2 |
| 389. | JUN | protein-protein interactions | ESR1 |
| 390. | JUN | protein-protein interactions | ESR2 |
| 391. | JUN | protein-protein interactions | RUNX2 |
| 392. | JUN | protein-protein interactions | estrogen receptor |
| 393. | JUNB | expression | RUNX2 |
| 394. | JUNB | protein-protein interactions | ESR1 |
| 395. | JUNB | protein-protein interactions | RUNX2 |
| 396. | JUND | protein-DNA interactions | RUNX2 |
| 397. | JUND | protein-protein interactions | ESR1 |
| 398. | KAT2B | protein-protein interactions | ESR1 |
| 399. | KAT2B | protein-protein interactions | estrogen receptor |
| 400. | KAT6A | expression | RUNX2 |
| 401. | KAT6A | protein-protein interactions | ESR1 |
| 402. | KAT6A | protein-protein interactions | RUNX2 |
| 403. | KAT6B | protein-protein interactions | ESR2 |
| 404. | KLF4 | expression | RUNX2 |
| 405. | KLF4 | protein-DNA interactions | RUNX2 |
| 406. | KLF4 | protein-protein interactions | ESR1 |
| 407. | KLF4 | protein-protein interactions | RUNX2 |
| 408. | LEF1 | expression | RUNX2 |
| 409. | LEF1 | inhibition | RUNX2 |
| 410. | LEF1 | protein-DNA interactions | RUNX2 |
| 411. | LEF1 | protein-protein interactions | ESR1 |
| 412. | LEF1 | protein-protein interactions | RUNX2 |
| 413. | LIMA1 | protein-protein interactions | ESR1 |
| 414. | LIMA1 | protein-protein interactions | HSP90 (family) |
| 415. | MAP2K1 | activation | RUNX2 |
| 416. | MAP2K1 | phosphorylation | RUNX2 |
| 417. | MAP2K1 | protein-protein interactions | ERBB2 |
| 418. | MAP2K1 | protein-protein interactions | HSP90 (family) |
| 419. | MATK | protein-protein interactions | ERBB2 |
| 420. | MEN1 | expression | RUNX2 |
| 421. | MEN1 | protein-protein interactions | ESR1 |
| 422. | MEN1 | protein-protein interactions | ESR2 |
| 423. | MEN1 | protein-protein interactions | RUNX2 |
| 424. | MEN1 | protein-protein interactions | estrogen receptor |
| 425. | MSX2 | activation | RUNX2 |
| 426. | MSX2 | protein-protein interactions | ESR1 |
| 427. | MSX2 | protein-protein interactions | RUNX2 |
| 428. | NFYB | protein-DNA interactions | RUNX2 |
| 429. | NFYB | protein-protein interactions | RUNX2 |
| 430. | NOTCH1 | activation | RUNX2 |
| 431. | NOTCH1 | expression | RUNX2 |

| | | | |
|---|---|---|---|
| 432. | NOTCH1 | protein-protein interactions | ERBB2 |
| 433. | NOTCH1 | protein-protein interactions | RUNX2 |
| 434. | NR0B2 | protein-protein interactions | ESR1 |
| 435. | NR0B2 | protein-protein interactions | ESR2 |
| 436. | NR0B2 | protein-protein interactions | estrogen receptor |
| 437. | NR3C1 | protein-DNA interactions | RUNX2 |
| 438. | NR3C1 | protein-protein interactions | ESR1 |
| 439. | NR3C1 | protein-protein interactions | FKBP4 |
| 440. | NR3C1 | protein-protein interactions | HSP90 (family) |
| 441. | NR3C1 | protein-protein interactions | estrogen receptor |
| 442. | OSTF1 | protein-DNA interactions | RUNX2 |
| 443. | PIN1 | activation | RUNX2 |
| 444. | PIN1 | protein-protein interactions | ERBB2 |
| 445. | PIN1 | protein-protein interactions | ESR1 |
| 446. | PIN1 | protein-protein interactions | RUNX2 |
| 447. | PML | protein-protein interactions | ESR2 |
| 448. | PPARD | expression | RUNX2 |
| 449. | PPARD | protein-protein interactions | HSP90 (family) |
| 450. | PPARG | expression | RUNX2 |
| 451. | PPARG | protein-protein interactions | ESR1 |
| 452. | PPARG | protein-protein interactions | HSP90 (family) |
| 453. | PPARG | protein-protein interactions | RUNX2 |
| 454. | PPARGC1A | protein-protein interactions | ESR1 |
| 455. | PPARGC1A | protein-protein interactions | ESR2 |
| 456. | PPARGC1A | transcription | RUNX2 |
| 457. | PPARGC1B | protein-protein interactions | ESR1 |
| 458. | PPARGC1B | transcription | RUNX2 |
| 459. | PTH | chemical-protein interactions | estrogen |
| 460. | RB | activation | RUNX2 |
| 461. | RB | protein-protein interactions | RUNX2 |
| 462. | RB1 | protein-protein interactions | ESR2 |
| 463. | RB1 | protein-protein interactions | estrogen receptor |
| 464. | RBM14 | protein-protein interactions | ERBB2 |
| 465. | RBM14 | protein-protein interactions | ESR1 |
| 466. | RBM28 | protein-protein interactions | ESR1 |
| 467. | RELA | expression | RUNX2 |
| 468. | RELA | protein-protein interactions | ESR1 |
| 469. | RUNX1 | expression | RUNX2 |
| 470. | RUNX1 | protein-DNA interactions | RUNX2 |
| 471. | RUNX1 | protein-protein interactions | ESR1 |
| 472. | RUNX1 | protein-protein interactions | RUNX2 |
| 473. | RUNX2 | activation | NOTCH1 |
| 474. | RUNX2 | activation | RUNX2 |
| 475. | RUNX2 | expression | RUNX2 |

| | | | |
|---|---|---|---|
| 476. | RUNX2 | inhibition | RUNX2 |
| 477. | RUNX2 | localization | RUNX2 |
| 478. | RUNX2 | modification | RUNX2 |
| 479. | RUNX2 | molecular cleavage | RUNX2 |
| 480. | RUNX2 | protein-DNA interactions | RUNX2 |
| 481. | RUNX2 | protein-protein interactions | ALYREF |
| 482. | RUNX2 | protein-protein interactions | AR |
| 483. | RUNX2 | protein-protein interactions | Ap1 |
| 484. | RUNX2 | protein-protein interactions | CCNB1 |
| 485. | RUNX2 | protein-protein interactions | CEBPB |
| 486. | RUNX2 | protein-protein interactions | CEBPD |
| 487. | RUNX2 | protein-protein interactions | CIC |
| 488. | RUNX2 | protein-protein interactions | CREBBP |
| 489. | RUNX2 | protein-protein interactions | CTBP2 |
| 490. | RUNX2 | protein-protein interactions | CTNNB1 |
| 491. | RUNX2 | protein-protein interactions | CUL4A |
| 492. | RUNX2 | protein-protein interactions | CUL4B |
| 493. | RUNX2 | protein-protein interactions | DDX5 |
| 494. | RUNX2 | protein-protein interactions | DET1 |
| 495. | RUNX2 | protein-protein interactions | EHMT2 |
| 496. | RUNX2 | protein-protein interactions | EP300 |
| 497. | RUNX2 | protein-protein interactions | ERK1/2 |
| 498. | RUNX2 | protein-protein interactions | ESR1 |
| 499. | RUNX2 | protein-protein interactions | ETS1 |
| 500. | RUNX2 | protein-protein interactions | EWSR1 |
| 501. | RUNX2 | protein-protein interactions | FHL2 |
| 502. | RUNX2 | protein-protein interactions | FOS |
| 503. | RUNX2 | protein-protein interactions | FOXO1 |
| 504. | RUNX2 | protein-protein interactions | GATA3 |
| 505. | RUNX2 | protein-protein interactions | GLI2 |
| 506. | RUNX2 | protein-protein interactions | GNB2 |
| 507. | RUNX2 | protein-protein interactions | GSN |
| 508. | RUNX2 | protein-protein interactions | HDAC1 |
| 509. | RUNX2 | protein-protein interactions | HDAC3 |
| 510. | RUNX2 | protein-protein interactions | HDAC4 |
| 511. | RUNX2 | protein-protein interactions | HDAC5 |
| 512. | RUNX2 | protein-protein interactions | HDAC6 |
| 513. | RUNX2 | protein-protein interactions | HDAC7 |
| 514. | RUNX2 | protein-protein interactions | HES1 |
| 515. | RUNX2 | protein-protein interactions | HEY1 |
| 516. | RUNX2 | protein-protein interactions | HIF1A |
| 517. | RUNX2 | protein-protein interactions | HMGB2 |
| 518. | RUNX2 | protein-protein interactions | HSPA4 |
| 519. | RUNX2 | protein-protein interactions | HSPA4L |

| | | | |
|---|---|---|---|
| 520. | RUNX2 | protein-protein interactions | HSPD1 |
| 521. | RUNX2 | protein-protein interactions | HSPH1 |
| 522. | RUNX2 | protein-protein interactions | IFI16 |
| 523. | RUNX2 | protein-protein interactions | IMPDH1 |
| 524. | RUNX2 | protein-protein interactions | JAG2 |
| 525. | RUNX2 | protein-protein interactions | JUN |
| 526. | RUNX2 | protein-protein interactions | JUNB |
| 527. | RUNX2 | protein-protein interactions | KAT2B |
| 528. | RUNX2 | protein-protein interactions | KAT6A |
| 529. | RUNX2 | protein-protein interactions | KAT6B |
| 530. | RUNX2 | protein-protein interactions | KLF4 |
| 531. | RUNX2 | protein-protein interactions | LEF1 |
| 532. | RUNX2 | protein-protein interactions | LIMA1 |
| 533. | RUNX2 | protein-protein interactions | MATK |
| 534. | RUNX2 | protein-protein interactions | MEN1 |
| 535. | RUNX2 | protein-protein interactions | MSX2 |
| 536. | RUNX2 | protein-protein interactions | NFYB |
| 537. | RUNX2 | protein-protein interactions | NOTCH1 |
| 538. | RUNX2 | protein-protein interactions | NR0B2 |
| 539. | RUNX2 | protein-protein interactions | PIN1 |
| 540. | RUNX2 | protein-protein interactions | PML |
| 541. | RUNX2 | protein-protein interactions | PPARG |
| 542. | RUNX2 | protein-protein interactions | RB |
| 543. | RUNX2 | protein-protein interactions | RB1 |
| 544. | RUNX2 | protein-protein interactions | RBL2 |
| 545. | RUNX2 | protein-protein interactions | RBM14 |
| 546. | RUNX2 | protein-protein interactions | RBM28 |
| 547. | RUNX2 | protein-protein interactions | RUNX1 |
| 548. | RUNX2 | protein-protein interactions | SMAD4 |
| 549. | RUNX2 | protein-protein interactions | SMAD5 |
| 550. | RUNX2 | protein-protein interactions | SNAI1 |
| 551. | RUNX2 | protein-protein interactions | osteocalcin |
| 552. | RUNX2 | regulation of binding | RUNX2 |
| 553. | RUNX2 | ubiquitination | RUNX2 |
| 554. | SIRT1 | expression | RUNX2 |
| 555. | SIRT1 | protein-protein interactions | AFP |
| 556. | SIRT1 | protein-protein interactions | ESR1 |
| 557. | SIRT1 | protein-protein interactions | RUNX2 |
| 558. | SKIC2 | protein-protein interactions | ESR1 |
| 559. | SKIC2 | protein-protein interactions | ESR2 |
| 560. | SKIC2 | protein-protein interactions | RUNX2 |
| 561. | SKP2 | protein-protein interactions | ESR1 |
| 562. | SKP2 | protein-protein interactions | FKBP4 |
| 563. | SKP2 | protein-protein interactions | RUNX2 |

| | | | |
|---|---|---|---|
| 564. | SMAD1 | expression | RUNX2 |
| 565. | SMAD1 | protein-protein interactions | ERBB2 |
| 566. | SMAD1 | protein-protein interactions | ESR1 |
| 567. | SMAD1 | protein-protein interactions | RUNX2 |
| 568. | SMAD2 | expression | RUNX2 |
| 569. | SMAD2 | protein-protein interactions | ESR1 |
| 570. | SMAD2 | protein-protein interactions | RUNX2 |
| 571. | SMAD2 | protein-protein interactions | estrogen receptor |
| 572. | SMAD3 | expression | RUNX2 |
| 573. | SMAD3 | protein-protein interactions | ESR1 |
| 574. | SMAD3 | protein-protein interactions | ESR2 |
| 575. | SMAD3 | protein-protein interactions | RUNX2 |
| 576. | SMAD3 | protein-protein interactions | estrogen receptor |
| 577. | SMAD4 | expression | RUNX2 |
| 578. | SMAD4 | protein-DNA interactions | RUNX2 |
| 579. | SMAD4 | protein-protein interactions | ESR1 |
| 580. | SMAD4 | protein-protein interactions | ESR2 |
| 581. | SMAD4 | protein-protein interactions | RUNX2 |
| 582. | SMAD4 | protein-protein interactions | estrogen receptor |
| 583. | SMAD5 | expression | RUNX2 |
| 584. | SMAD5 | protein-protein interactions | RUNX2 |
| 585. | SMAD6 | inhibition | RUNX2 |
| 586. | SMAD6 | protein-protein interactions | RUNX2 |
| 587. | SMARCA4 | expression | RUNX2 |
| 588. | SMARCA4 | protein-DNA interactions | RUNX2 |
| 589. | SMARCA4 | protein-protein interactions | ESR1 |
| 590. | SMARCA4 | protein-protein interactions | ESR2 |
| 591. | SMARCA4 | protein-protein interactions | RUNX2 |
| 592. | SMARCA4 | protein-protein interactions | estrogen receptor |
| 593. | SMURF1 | inhibition | RUNX2 |
| 594. | SMURF1 | protein-protein interactions | ESR1 |
| 595. | SMURF1 | protein-protein interactions | RUNX2 |
| 596. | SMURF1 | ubiquitination | RUNX2 |
| 597. | SMURF2 | protein-protein interactions | ERBB2 |
| 598. | SMURF2 | protein-protein interactions | FKBP4 |
| 599. | SMURF2 | protein-protein interactions | RUNX2 |
| 600. | SNAI1 | protein-protein interactions | RUNX2 |
| 601. | SNAI1 | transcription | RUNX2 |
| 602. | SNAI2 | activation | RUNX2 |
| 603. | SNAI2 | transcription | RUNX2 |
| 604. | SOX2 | protein-protein interactions | FKBP4 |
| 605. | SOX2 | protein-protein interactions | HSP90 (family) |
| 606. | SOX2 | protein-protein interactions | RUNX2 |
| 607. | SOX9 | expression | RUNX2 |

| | | | |
|---|---|---|---|
| 608. | SOX9 | protein-protein interactions | RUNX2 |
| 609. | SP1 | expression | ESR1 |
| 610. | SP1 | expression | RUNX2 |
| 611. | SP1 | protein-DNA interactions | RUNX2 |
| 612. | SP1 | protein-protein interactions | ESR1 |
| 613. | SP1 | protein-protein interactions | ESR2 |
| 614. | SP1 | protein-protein interactions | estrogen receptor |
| 615. | SP1 | transcription | RUNX2 |
| 616. | SRF | activation | RUNX2 |
| 617. | SRF | protein-protein interactions | RUNX2 |
| 618. | STAT1 | inhibition | RUNX2 |
| 619. | STAT1 | protein-protein interactions | ERBB2 |
| 620. | STAT1 | protein-protein interactions | ESR1 |
| 621. | STAT1 | protein-protein interactions | ESR2 |
| 622. | STAT1 | protein-protein interactions | RUNX2 |
| 623. | STAT3 | protein-protein interactions | ERBB2 |
| 624. | STAT3 | protein-protein interactions | ESR1 |
| 625. | STAT3 | protein-protein interactions | ESR2 |
| 626. | STAT3 | protein-protein interactions | HSP90 (family) |
| 627. | STAT3 | protein-protein interactions | RUNX2 |
| 628. | STAT5A | expression | RUNX2 |
| 629. | STAT5A | protein-protein interactions | ESR1 |
| 630. | STAT5A | protein-protein interactions | ESR2 |
| 631. | STAT5A | protein-protein interactions | RUNX2 |
| 632. | STUB1 | protein-protein interactions | ERBB2 |
| 633. | STUB1 | protein-protein interactions | ESR1 |
| 634. | STUB1 | protein-protein interactions | ESR2 |
| 635. | STUB1 | protein-protein interactions | HSP90 (family) |
| 636. | STUB1 | protein-protein interactions | RUNX2 |
| 637. | SUV39H1 | protein-protein interactions | ESR1 |
| 638. | SUV39H1 | protein-protein interactions | RUNX2 |
| 639. | TAF1A | protein-protein interactions | ESR1 |
| 640. | TAF1A | protein-protein interactions | RUNX2 |
| 641. | TCF7L2 | protein-protein interactions | ESR1 |
| 642. | TCF7L2 | protein-protein interactions | ESR2 |
| 643. | TCF7L2 | protein-protein interactions | RUNX2 |
| 644. | TFAM | expression | RUNX2 |
| 645. | TFAM | protein-protein interactions | ESR1 |
| 646. | THRAP3 | expression | RUNX2 |
| 647. | THRAP3 | protein-protein interactions | ESR1 |
| 648. | THRB | expression | RUNX2 |
| 649. | THRB | protein-DNA interactions | RUNX2 |
| 650. | THRB | protein-protein interactions | estrogen receptor |
| 651. | THRB | transcription | RUNX2 |

| | | | |
|---|---|---|---|
| 652. | TLE1 | activation | RUNX2 |
| 653. | TLE1 | protein-protein interactions | ESR1 |
| 654. | TLE1 | protein-protein interactions | RUNX2 |
| 655. | TP53 | expression | ESR1 |
| 656. | TP53 | expression | RUNX2 |
| 657. | TP53 | protein-protein interactions | ESR1 |
| 658. | TP53 | protein-protein interactions | ESR2 |
| 659. | TP53 | protein-protein interactions | FKBP4 |
| 660. | TP53 | protein-protein interactions | HSP90 (family) |
| 661. | TP53 | protein-protein interactions | PTH |
| 662. | TP53 | protein-protein interactions | RUNX2 |
| 663. | TP53 | transcription | RUNX2 |
| 664. | TP73 | protein-protein interactions | HSP90 (family) |
| 665. | TP73 | protein-protein interactions | RUNX2 |
| 666. | TRIB3 | expression | RUNX2 |
| 667. | TRPS1 | expression | RUNX2 |
| 668. | TRPS1 | protein-protein interactions | ESR1 |
| 669. | TSC22D3 | expression | RUNX2 |
| 670. | TWIST1 | expression | RUNX2 |
| 671. | TWIST1 | protein-protein interactions | RUNX2 |
| 672. | UBTF | protein-protein interactions | ESR1 |
| 673. | UBTF | protein-protein interactions | RUNX2 |
| 674. | VDR | expression | RUNX2 |
| 675. | VDR | protein-protein interactions | HSP90 (family) |
| 676. | VDR | protein-protein interactions | RUNX2 |
| 677. | WDR5 | expression | RUNX2 |
| 678. | WDR5 | protein-DNA interactions | RUNX2 |
| 679. | WDR5 | protein-protein interactions | ESR1 |
| 680. | WDR5 | protein-protein interactions | ESR2 |
| 681. | WWP1 | protein-protein interactions | ESR2 |
| 682. | WWP1 | protein-protein interactions | RUNX2 |
| 683. | WWP1 | ubiquitination | RUNX2 |
| 684. | WWP2 | protein-protein interactions | ERBB2 |
| 685. | WWP2 | protein-protein interactions | RUNX2 |
| 686. | XRCC5 | protein-protein interactions | ESR1 |
| 687. | XRCC5 | protein-protein interactions | ESR2 |
| 688. | XRCC5 | protein-protein interactions | RUNX2 |
| 689. | XRCC6 | protein-protein interactions | ESR1 |
| 690. | XRCC6 | protein-protein interactions | RUNX2 |
| 691. | YAP1 | expression | RUNX2 |
| 692. | YAP1 | protein-protein interactions | ESR2 |
| 693. | YAP1 | protein-protein interactions | RUNX2 |
| 694. | YY1 | activation | RUNX2 |
| 695. | YY1 | protein-protein interactions | RUNX2 |

| | | | |
|---|---|---|---|
| 696. | ZBTB16 | expression | RUNX2 |
| 697. | ZBTB16 | protein-protein interactions | ESR1 |
| 698. | ZBTB7B | protein-protein interactions | ESR1 |
| 699. | ZBTB7B | protein-protein interactions | ESR2 |
| 700. | ZBTB7B | protein-protein interactions | RUNX2 |
| 701. | ZMYND8 | expression | RUNX2 |
| 702. | ZMYND8 | protein-protein interactions | ERBB2 |
| 703. | ZMYND8 | protein-protein interactions | ESR1 |
| 704. | ZMYND8 | protein-protein interactions | ESR2 |
| 705. | estrogen | activation | ERBB2 |
| 706. | estrogen | activation | ESR1 |
| 707. | estrogen | activation | ESR2 |
| 708. | estrogen | activation | RUNX2 |
| 709. | estrogen | activation | estrogen receptor |
| 710. | estrogen | chemical-protein interactions | AFP |
| 711. | estrogen | chemical-protein interactions | ERBB2 |
| 712. | estrogen | chemical-protein interactions | ESR1 |
| 713. | estrogen | chemical-protein interactions | ESR2 |
| 714. | estrogen | chemical-protein interactions | estrogen receptor |
| 715. | estrogen | inhibition | ERBB2 |
| 716. | estrogen | reaction | ER/estrogen |
| 717. | estrogen | reaction | ESTG:ESR1:chaperone |
| 718. | estrogen | reaction | ESTG:ESR2:chaperone |
| 719. | estrogen | reaction | ESTG:Me-PalmS-ESR dimers |
| 720. | estrogen | translocation | estrogen |
| 721. | estrogen receptor | activation | FOS |
| 722. | estrogen receptor | activation | JUN |
| 723. | estrogen receptor | activation | TP53 |
| 724. | estrogen receptor | chemical-protein interactions | estrogen |
| 725. | estrogen receptor | inhibition | DDX5 |
| 726. | estrogen receptor | membership | ESR1 |
| 727. | estrogen receptor | membership | ESR2 |
| 728. | estrogen receptor | protein-DNA interactions | FOS |
| 729. | estrogen receptor | protein-protein interactions | AR |
| 730. | estrogen receptor | protein-protein interactions | CREBBP |
| 731. | estrogen receptor | protein-protein interactions | DDX5 |
| 732. | estrogen receptor | protein-protein interactions | EP300 |
| 733. | estrogen receptor | protein-protein interactions | FOS |
| 734. | estrogen receptor | protein-protein interactions | JUN |
| 735. | histone H3 | protein-DNA interactions | RUNX2 |
| 736. | histone H4 | protein-DNA interactions | RUNX2 |

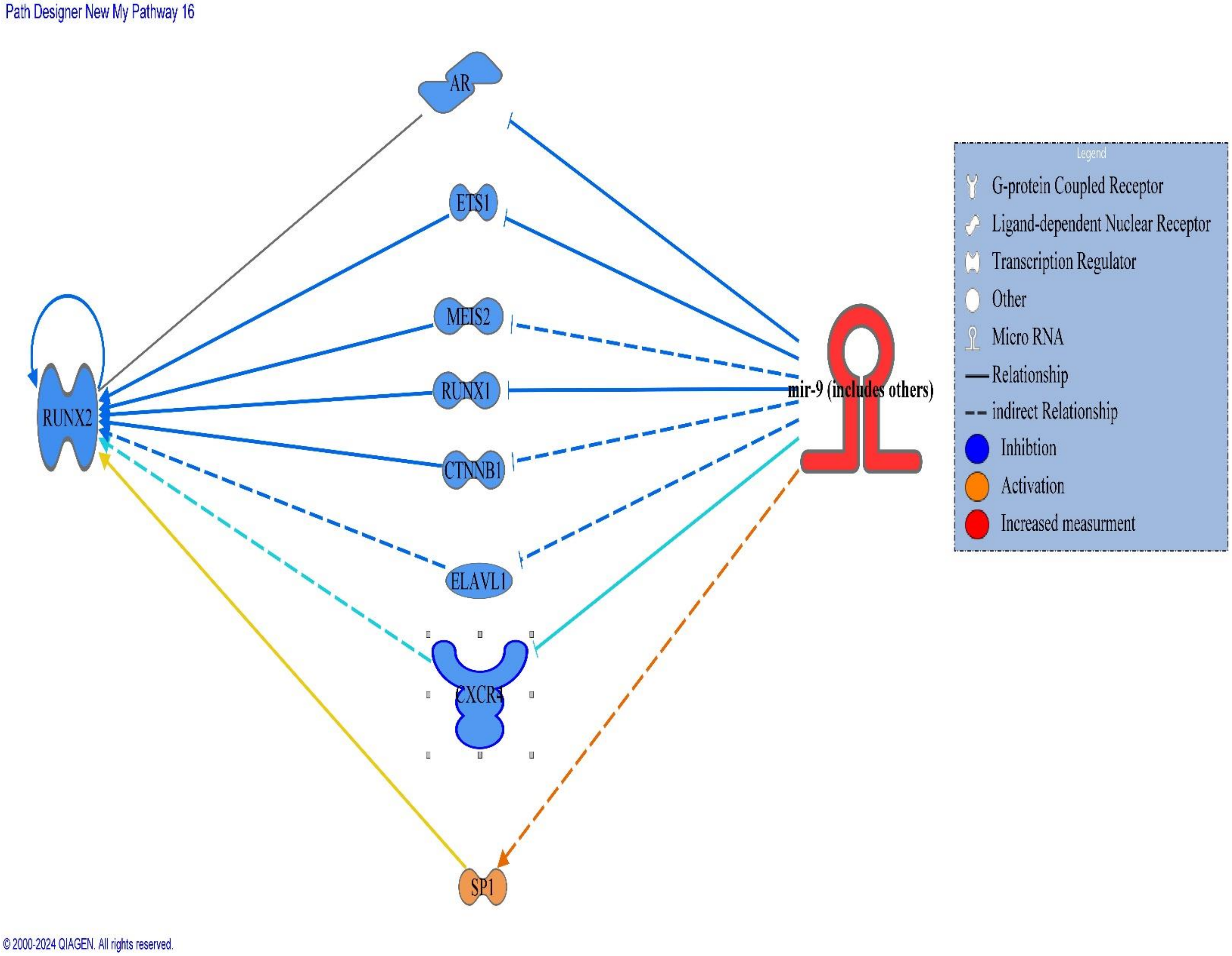


Figure.5: Molecular network depicting the connectivity and relationships among the overlapping molecules associated with direct influence of miRNA9 on RUNX-2 expression

Table.5: Various molecules incorporated in the miRNA-9 and *RUNX-2 gene* expression

| Symbol | Molecule/ Gene Name | Location | Family |
|---|---|---|---|
| 1. AR | androgen receptor | Nucleus | ligand-dependent nuclear receptor |
| 2. CTNNB1 | catenin beta 1 | Nucleus | transcription regulator |
| 3. CXCR4 | C-X-C motif chemokine receptor 4 | Plasma Membrane | G-protein coupled receptor |
| 4. ELAVL1 | ELAV like RNA binding protein 1 | Cytoplasm | other |
| 5. ETS1 | ETS proto-oncogene 1, transcription factor | Nucleus | transcription regulator |
| 6. MEIS2 | Meis homeobox 2 | Nucleus | transcription regulator |
| 7. mir-9 (includes others) | relatives of microRNA 9 | Cytoplasm | microRNA |
| 8. RUNX1 | RUNX family transcription factor 1 | Nucleus | transcription regulator |
| 9. RUNX2 | RUNX family transcription factor 2 | Nucleus | transcription regulator |
| 10. SP1 | Sp1 transcription factor | Nucleus | transcription regulator |

Table.6: Various interactions between molecules incorporated in the miRNA-9 and *RUNX-2 gene* expression

| From Molecule(s) | Relationship Type | To Molecule(s) |
|---|---|---|
| 1. CTNNB1 | expression | RUNX2 |
| 2. CTNNB1 | inhibition | RUNX2 |
| 3. CTNNB1 | protein-DNA interactions | RUNX2 |
| 4. CTNNB1 | protein-protein interactions | RUNX2 |
| 5. CTNNB1 | transcription | RUNX2 |
| 6. CXCR4 | expression | RUNX2 |
| 7. ELAVL1 | expression | RUNX2 |
| 8. ETS1 | protein-DNA interactions | RUNX2 |
| 9. ETS1 | protein-protein interactions | RUNX2 |
| 10. ETS1 | transcription | RUNX2 |
| 11. MEIS2 | expression | RUNX2 |
| 12. RUNX1 | expression | RUNX2 |
| 13. RUNX1 | protein-DNA interactions | RUNX2 |
| 14. RUNX1 | protein-protein interactions | RUNX2 |
| 15. RUNX2 | activation | RUNX2 |
| 16. RUNX2 | expression | RUNX2 |
| 17. RUNX2 | inhibition | RUNX2 |
| 18. RUNX2 | localization | RUNX2 |
| 19. RUNX2 | modification | RUNX2 |
| 20. RUNX2 | molecular cleavage | RUNX2 |
| 21. RUNX2 | protein-DNA interactions | RUNX2 |
| 22. RUNX2 | protein-protein interactions | AR |
| 23. RUNX2 | protein-protein interactions | CTNNB1 |
| 24. RUNX2 | protein-protein interactions | ETS1 |
| 25. RUNX2 | protein-protein interactions | RUNX1 |
| 26. RUNX2 | regulation of binding | RUNX2 |
| 27. RUNX2 | ubiquitination | RUNX2 |
| 28. SP1 | expression | RUNX2 |

| | | |
|---|---|---|
| 29.SP1 | protein-DNA interactions | RUNX2 |
| 30.SP1 | transcription | RUNX2 |
| 31.mir-9 | RNA-RNA interactions: microRNA targeting | AR |
| 32.mir-9 | RNA-RNA interactions: microRNA targeting | CXCR4 |
| 33.mir-9 | RNA-RNA interactions: microRNA targeting | ETS1 |
| 34.mir-9 | RNA-RNA interactions: microRNA targeting | RUNX1 |
| 35.mir-9 | RNA-RNA interactions: non-targeting interactions | AR |
| 36.mir-9 | expression | AR |
| 37.mir-9 | expression | CTNNB1 |
| 38.mir-9 | expression | CXCR4 |
| 39.mir-9 | expression | ETS1 |
| 40.mir-9 | expression | MEIS2 |
| 41.mir-9 | expression | RUNX1 |
| 42.mir-9 | expression | SP1 |
| 43.mir-9 | inhibition | ELAVL1 |

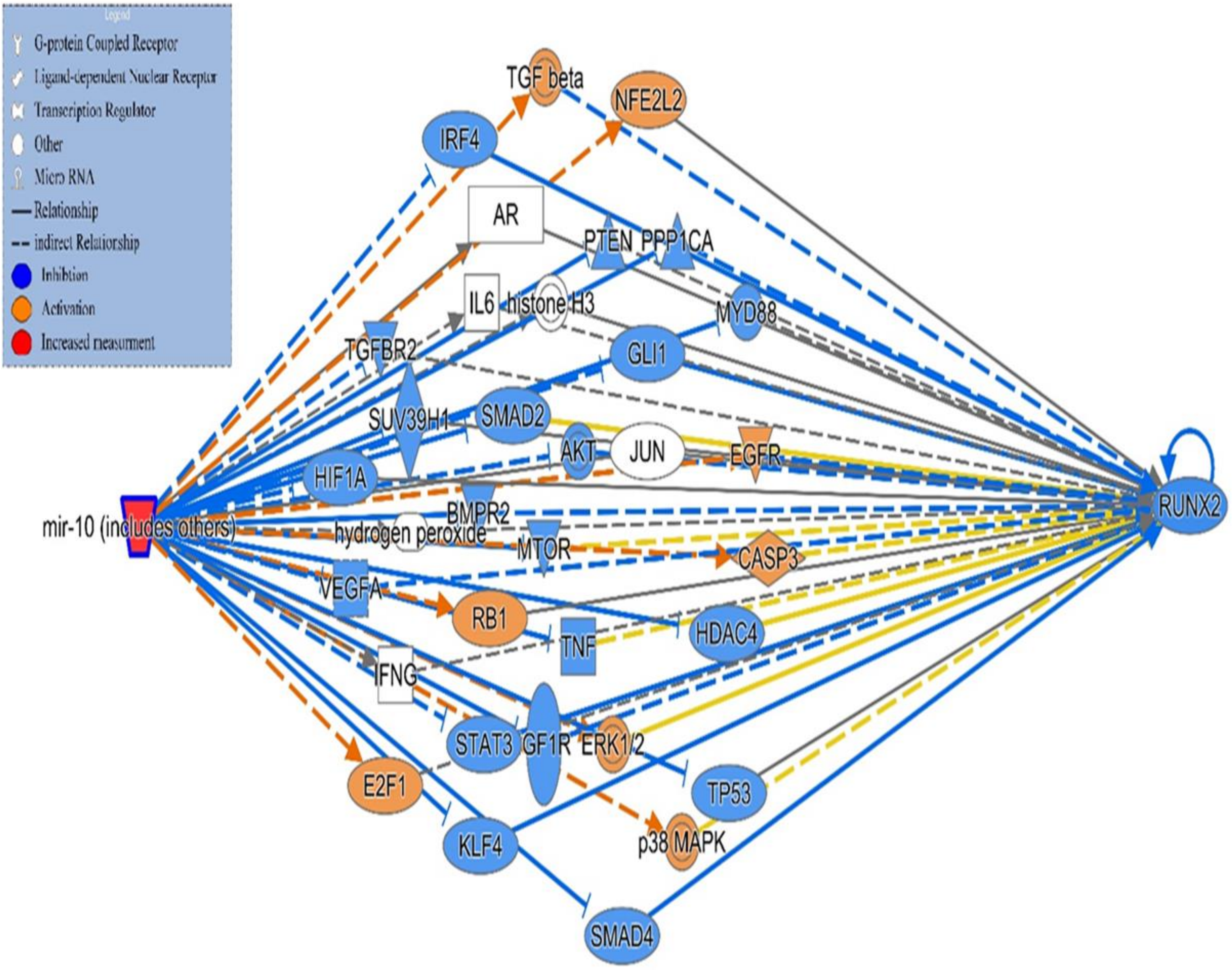


Figure.6: Molecular network depicting the connectivity and relationships among the overlapping molecules associated with direct influence of miRNA10 on *RUNX-2* expression

**Table(7): Various molecules incorporated in the regulation of miRNA-10 and affect *RUNX-2 gene* expression**

| Symbol | Molecule/ Gene Name | Location | Family |
|---|---|---|---|
| 1. AKT | | Cytoplasm | group |
| 2. AR | androgen receptor | Nucleus | ligand-dependent nuclear receptor |
| 3. BMPR2 | bone morphogenetic protein receptor type 2 | Plasma Membrane | kinase |
| 4. CASP3 | caspase 3 | Cytoplasm | peptidase |
| 5. E2F1 | E2F transcription factor 1 | Nucleus | transcription regulator |
| 6. EGFR | epidermal growth factor receptor | Plasma Membrane | kinase |
| 7. ERK1/2 | | Cytoplasm | group |
| 8. GLI1 | GLI family zinc finger 1 | Nucleus | transcription regulator |
| 9. HDAC4 | histone deacetylase 4 | Nucleus | transcription regulator |

| | | | |
|---|---|---|---|
| 10. HIF1A | hypoxia inducible factor 1 subunit alpha | Nucleus | transcription regulator |
| 11. histone H3 | | Nucleus | Group |
| 12. hydrogen peroxide | | Other | chemical - endogenous mammalian |
| 13. IFNG | interferon gamma | Extracellular Space | Cytokine |
| 14. IGF1R | insulin like growth factor 1 receptor | Plasma Membrane | transmembrane receptor |
| 15. IL6 | interleukin 6 | Extracellular Space | Cytokine |
| 16. IRF4 | interferon regulatory factor 4 | Nucleus | transcription regulator |
| 17. JUN | Jun proto-oncogene, AP-1 transcription factor subunit | Nucleus | transcription regulator |
| 18. KLF4 | KLF transcription factor 4 | Nucleus | transcription regulator |
| 19. mir-10 (includes others) | relatives of microRNA 10 | Cytoplasm | microRNA |
| 20. MTOR | mechanistic target of rapamycin kinase | Nucleus | Kinase |
| 21. MYD88 | MYD88 innate immune signal transduction adaptor | Plasma Membrane | other |
| 22. NFE2L2 | NFE2 like bZIP transcription factor 2 | Nucleus | transcription regulator |
| 23. p38 MAPK | | Cytoplasm | group |
| 24. PPP1CA | protein phosphatase 1 catalytic subunit alpha | Cytoplasm | phosphatase |
| 25. PTEN | phosphatase and tensin homolog | Cytoplasm | phosphatase |
| 26. RB1 | RB transcriptional corepressor 1 | Nucleus | transcription regulator |
| 27. RUNX2 | RUNX family transcription factor 2 | Nucleus | transcription regulator |
| 28. SMAD2 | SMAD family member 2 | Nucleus | transcription regulator |
| 29. SMAD4 | SMAD family member 4 | Nucleus | transcription regulator |
| 30. STAT3 | signal transducer and activator of transcription 3 | Nucleus | transcription regulator |
| 31. SUV39H1 | SUV39H1 histone lysine methyltransferase | Nucleus | enzyme |
| 32. TGF beta | | Extracellular Space | group |
| 33. TGFBR2 | transforming growth factor beta receptor 2 | Plasma Membrane | kinase |
| 34. TNF | tumor necrosis factor | Extracellular Space | cytokine |
| 35. TP53 | tumor protein p53 | Nucleus | transcription regulator |
| 36. VEGFA | vascular endothelial growth factor A | Extracellular Space | growth factor |

**Table.8: Various interactions between molecules incorporated in the miRNA-10 and RUNX-2 gene expression**

| From Molecule(s) | Relationship Type | To Molecule(s) |
|---|---|---|
| 1. AKT | activation | RUNX2 |
| 2. AKT | expression | RUNX2 |
| 3. AKT | phosphorylation | RUNX2 |
| 4. BMPR2 | expression | RUNX2 |
| 5. BMPR2 | regulation of binding | RUNX2 |
| 6. CASP3 | expression | RUNX2 |
| 7. E2F1 | regulation of binding | RUNX2 |
| 8. EGFR | expression | RUNX2 |
| 9. ERK1/2 | activation | RUNX2 |
| 10. ERK1/2 | expression | RUNX2 |
| 11. ERK1/2 | phosphorylation | RUNX2 |
| 12. ERK1/2 | protein-protein interactions | RUNX2 |
| 13. GLI1 | expression | RUNX2 |
| 14. GLI1 | protein-protein interactions | RUNX2 |
| 15. GLI1 | transcription | RUNX2 |
| 16. HDAC4 | activation | RUNX2 |
| 17. HDAC4 | expression | RUNX2 |
| 18. HDAC4 | modification | RUNX2 |
| 19. HDAC4 | protein-protein interactions | RUNX2 |
| 20. IFNG | expression | RUNX2 |
| 21. IGF1R | expression | RUNX2 |
| 22. IL6 | expression | RUNX2 |
| 23. IRF4 | expression | RUNX2 |
| 24. JUN | protein-DNA interactions | RUNX2 |
| 25. JUN | protein-protein interactions | RUNX2 |
| 26. KLF4 | expression | RUNX2 |
| 27. KLF4 | protein-DNA interactions | RUNX2 |
| 28. KLF4 | protein-protein interactions | RUNX2 |
| 29. MTOR | expression | RUNX2 |
| 30. MYD88 | expression | RUNX2 |
| 31. NFE2L2 | protein-protein interactions | RUNX2 |
| 32. NFE2L2 | regulation of binding | RUNX2 |
| 33. PPP1CA | expression | RUNX2 |
| 34. PTEN | regulation of binding | RUNX2 |
| 35. RUNX2 | activation | RUNX2 |
| 36. RUNX2 | expression | RUNX2 |
| 37. RUNX2 | inhibition | RUNX2 |
| 38. RUNX2 | localization | RUNX2 |
| 39. RUNX2 | modification | RUNX2 |
| 40. RUNX2 | molecular cleavage | RUNX2 |
| 41. RUNX2 | protein-DNA interactions | RUNX2 |
| 42. RUNX2 | protein-protein interactions | AR |
| 43. RUNX2 | protein-protein interactions | ERK1/2 |
| 44. RUNX2 | protein-protein interactions | GLI1 |
| 45. RUNX2 | protein-protein interactions | HDAC4 |
| 46. RUNX2 | protein-protein interactions | HIF1A |
| 47. RUNX2 | protein-protein interactions | JUN |
| 48. RUNX2 | protein-protein interactions | KLF4 |
| 49. RUNX2 | protein-protein interactions | NFE2L2 |
| 50. RUNX2 | protein-protein interactions | RB1 |
| 51. RUNX2 | protein-protein interactions | SMAD4 |

| | | |
|---|---|---|
| 52. RUNX2 | regulation of binding | RUNX2 |
| 53. RUNX2 | ubiquitination | RUNX2 |
| 54. SMAD2 | expression | RUNX2 |
| 55. SMAD2 | protein-protein interactions | RUNX2 |
| 56. SMAD4 | expression | RUNX2 |
| 57. SMAD4 | protein-DNA interactions | RUNX2 |
| 58. SMAD4 | protein-protein interactions | RUNX2 |
| 59. STAT3 | expression | RUNX2 |
| 60. STAT3 | protein-protein interactions | RUNX2 |
| 61. SUV39H1 | protein-protein interactions | RUNX2 |
| 62. SUV39H1 | protein-protein interactions | mir-10 (includes others) |
| 63. TGF beta | expression | RUNX2 |
| 64. TGFBR2 | expression | RUNX2 |
| 65. TNF | expression | RUNX2 |
| 66. TNF | localization | RUNX2 |
| 67. TNF | molecular cleavage | RUNX2 |
| 68. TNF | transcription | RUNX2 |
| 69. TP53 | expression | RUNX2 |
| 70. TP53 | protein-protein interactions | RUNX2 |
| 71. TP53 | transcription | RUNX2 |
| 72. VEGFA | expression | RUNX2 |
| 73. histone H3 | protein-DNA interactions | RUNX2 |
| 74. hydrogen peroxide | expression | RUNX2 |
| 75. mir-10 (includes others) | RNA-RNA interactions: microRNA targeting | AR |
| 76. mir-10 (includes others) | RNA-RNA interactions: microRNA targeting | BMPR2 |
| 77. mir-10 (includes others) | RNA-RNA interactions: microRNA targeting | HDAC4 |
| 78. mir-10 (includes others) | RNA-RNA interactions: microRNA targeting | IGF1R |
| 79. mir-10 (includes others) | RNA-RNA interactions: microRNA targeting | KLF4 |
| 80. mir-10 (includes others) | RNA-RNA interactions: microRNA targeting | MTOR |
| 81. mir-10 (includes others) | RNA-RNA interactions: microRNA targeting | MYD88 |
| 82. mir-10 (includes others) | RNA-RNA interactions: microRNA targeting | PPP1CA |
| 83. mir-10 (includes others) | RNA-RNA interactions: microRNA targeting | PTEN |
| 84. mir-10 (includes others) | RNA-RNA interactions: microRNA targeting | SMAD2 |
| 85. mir-10 (includes others) | RNA-RNA interactions: microRNA targeting | SMAD4 |
| 86. mir-10 (includes others) | RNA-RNA interactions: microRNA targeting | SUV39H1 |
| 87. mir-10 (includes others) | RNA-RNA interactions: microRNA targeting | TNF |
| 88. mir-10 (includes others) | RNA-RNA interactions: microRNA targeting | TP53 |
| 89. mir-10 (includes others) | RNA-RNA interactions: non-targeting interactions | JUN |
| 90. mir-10 (includes others) | activation | AKT |
| 91. mir-10 (includes others) | activation | CASP3 |
| 92. mir-10 (includes others) | activation | EGFR |
| 93. mir-10 (includes others) | activation | ERK1/2 |
| 94. mir-10 (includes others) | activation | STAT3 |
| 95. mir-10 (includes others) | activation | TP53 |
| 96. mir-10 (includes others) | expression | AKT |
| 97. mir-10 (includes others) | expression | AR |
| 98. mir-10 (includes others) | expression | BMPR2 |
| 99. mir-10 (includes others) | expression | E2F1 |
| 100. mir-10 (includes others) | expression | GLI1 |
| 101. mir-10 (includes others) | expression | HDAC4 |
| 102. mir-10 (includes others) | expression | HIF1A |
| 103. mir-10 (includes others) | expression | IGF1R |
| 104. mir-10 (includes others) | expression | IL6 |

| | | |
|---|---|---|
| 105.mir-10 (includes others) | expression | IRF4 |
| 106.mir-10 (includes others) | expression | KLF4 |
| 107.mir-10 (includes others) | expression | MTOR |
| 108.mir-10 (includes others) | expression | MYD88 |
| 109.mir-10 (includes others) | expression | NFE2L2 |
| 110.mir-10 (includes others) | expression | PPP1CA |
| 111.mir-10 (includes others) | expression | PTEN |
| 112.mir-10 (includes others) | expression | RB1 |
| 113.mir-10 (includes others) | expression | SMAD2 |
| 114.mir-10 (includes others) | expression | SMAD4 |
| 115.mir-10 (includes others) | expression | SUV39H1 |
| 116.mir-10 (includes others) | expression | TGF beta |
| 117.mir-10 (includes others) | expression | TGFBR2 |
| 118.mir-10 (includes others) | expression | TNF |
| 119.mir-10 (includes others) | expression | TP53 |
| 120.mir-10 (includes others) | expression | VEGFA |
| 121.mir-10 (includes others) | expression | p38 MAPK |
| 122.mir-10 (includes others) | localization | IFNG |
| 123.mir-10 (includes others) | localization | hydrogen peroxide |
| 124.mir-10 (includes others) | molecular cleavage | CASP3 |
| 125.mir-10 (includes others) | phosphorylation | AKT |
| 126.mir-10 (includes others) | phosphorylation | EGFR |
| 127.mir-10 (includes others) | phosphorylation | ERK1/2 |
| 128.mir-10 (includes others) | phosphorylation | STAT3 |
| 129.mir-10 (includes others) | protein-protein interactions | SUV39H1 |
| 130.mir-10 (includes others) | regulation of binding | IL6 |
| 131.mir-10 (includes others) | regulation of binding | histone H3 |
| 132.mir-10 (includes others) | translocation | STAT3 |
| 133.p38 MAPK | activation | RUNX2 |
| 134.p38 MAPK | expression | RUNX2 |

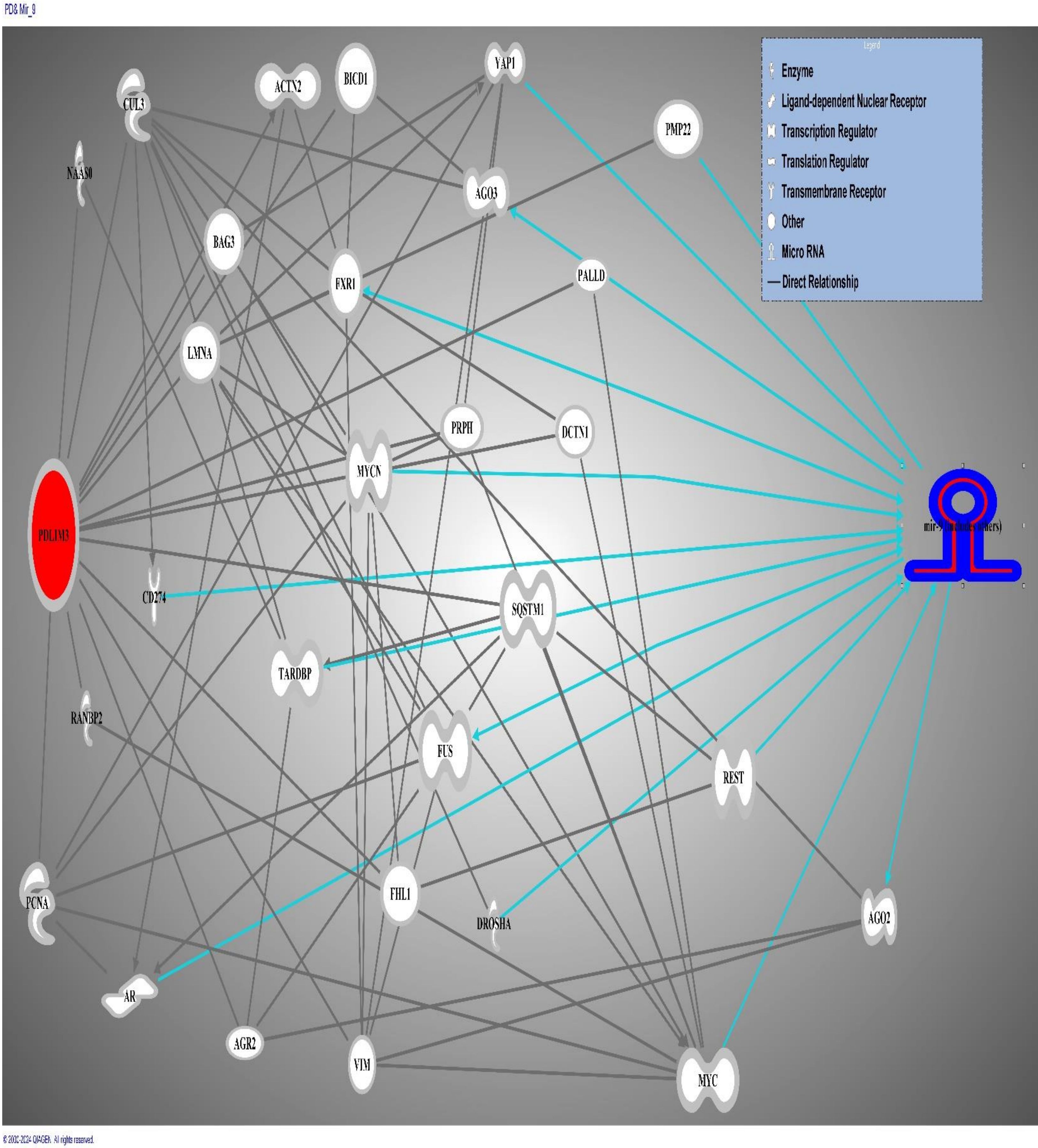


Figure.7: Molecular network depicting the connectivity and relationships among the overlapping molecules associated with direct influence of PDLIM-3 on miRNA-9 expression

**Table 9: Various molecules incorporated in the regulation of *PDLIM-3* and affect *miRNA9* expression**

| Symbol | Molecule/ Gene Name | Location | Family |
|---|---|---|---|
| 1. ACTN2 | actinin alpha 2 | Nucleus | transcription regulator |
| 2. AGO2 | argonaute RISC catalytic component 2 | Cytoplasm | translation regulator |
| 3. AGO3 | argonaute RISC catalytic component 3 | Cytoplasm | translation regulator |
| 4. AGR2 | anterior gradient 2, protein disulphide isomerase family member | Extracellular Space | other |
| 5. AR | androgen receptor | Nucleus | ligand-dependent nuclear receptor |
| 6. BAG3 | BAG cochaperone 3 | Cytoplasm | other |
| 7. BICD1 | BICD cargo adaptor 1 | Cytoplasm | other |
| 8. CD274 | CD274 molecule | Plasma Membrane | transmembrane receptor |
| 9. CUL3 | cullin 3 | Nucleus | enzyme |
| 10. DCTN1 | dynactin subunit 1 | Cytoplasm | other |
| 11. DROSHA | drosha ribonuclease III | Nucleus | enzyme |
| 12. FHL1 | four and a half LIM domains 1 | Cytoplasm | other |
| 13. FUS | FUS RNA binding protein | Nucleus | transcription regulator |
| 14. FXR1 | FMR1 autosomal homolog 1 | Cytoplasm | other |
| 15. LMNA | lamin A/C | Nucleus | other |
| 16. mir-9 | relatives of microRNA 9 | Cytoplasm | microRNA |
| 17. MYC | MYC proto-oncogene, bHLH transcription factor | Nucleus | transcription regulator |
| 18. MYCN | MYCN proto-oncogene, bHLH transcription factor | Nucleus | transcription regulator |
| 19. NAA40 | N-alpha-acetyltransferase 40, NatD catalytic subunit | Cytoplasm | enzyme |
| 20. PALLD | palladin, cytoskeletal associated protein | Plasma Membrane | other |
| 21. PCNA | proliferating cell nuclear antigen | Nucleus | enzyme |
| 22. PDLIM3 | PDZ and LIM domain 3 | Cytoplasm | other |
| 23. PMP22 | peripheral myelin protein 22 | Plasma Membrane | other |
| 24. PRPH | peripherin | Plasma Membrane | other |
| 25. RANBP2 | RAN binding protein 2 | Nucleus | enzyme |
| 26. REST | RE1 silencing transcription factor | Nucleus | transcription regulator |
| 27. SQSTM1 | sequestosome 1 | Cytoplasm | transcription regulator |
| 28. TARDBP | TAR DNA binding protein | Nucleus | transcription regulator |
| 29. VIM | vimentin | Cytoplasm | other |
| 30. YAP1 | Yes1 associated transcriptional regulator | Nucleus | transcription regulator |

**Table.10: Various interactions between molecules incorporated in the PDLIM-3 and affect *miRNA9* expression**

| From Molecule(s) | Relationship Type | To Molecule(s) |
|---|---|---|
| 1.ACTN2 | activation | AR |
| 2.ACTN2 | protein-protein interactions | AR |
| 3.AGR2 | protein-protein interactions | AGO2 |
| 4.AR | protein-protein interactions | ACTN2 |
| 5.BAG3 | expression | MYC |
| 6.BAG3 | protein-protein interactions | MYC |
| 7.BICD1 | protein-protein interactions | AGO3 |
| 8.CD274 | expression | mir-9 (includes others) |
| 9.CUL3 | expression | CD274 |
| 10. CUL3 | protein-protein interactions | AGO2 |
| 11. CUL3 | protein-protein interactions | AGO3 |
| 12. CUL3 | protein-protein interactions | CD274 |
| 13. DROSHA | protein-DNA interactions | mir-9 (includes others) |
| 14. DROSHA | protein-protein interactions | CUL3 |
| 15. FUS | protein-RNA interactions | mir-9 (includes others) |
| 16. FUS | protein-protein interactions | AGR2 |
| 17. FUS | protein-protein interactions | CUL3 |
| 18. FUS | regulation of binding | mir-9 (includes others) |
| 19. FXR1 | expression | mir-9 (includes others) |
| 20. FXR1 | molecular cleavage | mir-9 (includes others) |
| 21. FXR1 | protein-RNA interactions | mir-9 (includes others) |
| 22. FXR1 | protein-protein interactions | ACTN2 |
| 23. FXR1 | protein-protein interactions | BICD1 |
| 24. FXR1 | protein-protein interactions | CUL3 |
| 25. FXR1 | protein-protein interactions | DCTN1 |
| 26. LMNA | expression | MYC |
| 27. LMNA | inhibition | YAP1 |
| 28. LMNA | localization | YAP1 |
| 29. LMNA | phosphorylation | YAP1 |
| 30. LMNA | protein-protein interactions | FUS |
| 31. LMNA | protein-protein interactions | FXR1 |
| 32. LMNA | protein-protein interactions | MYC |
| 33. LMNA | protein-protein interactions | YAP1 |
| 34. LMNA | transcription | YAP1 |
| 35. MYC | expression | mir-9 (includes others) |

| | | | |
|---|---|---|---|
| 36. | MYC | protein-DNA interactions | mir-9 (includes others) |
| 37. | MYC | protein-protein interactions | BAG3 |
| 38. | MYC | protein-protein interactions | DCTN1 |
| 39. | MYC | protein-protein interactions | LMNA |
| 40. | MYC | transcription | mir-9 (includes others) |
| 41. | MYCN | protein-DNA interactions | mir-9 (includes others) |
| 42. | MYCN | protein-protein interactions | CUL3 |
| 43. | MYCN | protein-protein interactions | FHL1 |
| 44. | MYCN | protein-protein interactions | LMNA |
| 45. | PALLD | protein-protein interactions | MYC |
| 46. | PCNA | protein-protein interactions | AR |
| 47. | PCNA | protein-protein interactions | FUS |
| 48. | PCNA | protein-protein interactions | MYC |
| 49. | PCNA | protein-protein interactions | MYCN |
| 50. | PDLIM3 | protein-protein interactions | ACTN2 |
| 51. | PDLIM3 | protein-protein interactions | AGR2 |
| 52. | PDLIM3 | protein-protein interactions | BAG3 |
| 53. | PDLIM3 | protein-protein interactions | BICD1 |
| 54. | PDLIM3 | protein-protein interactions | CUL3 |
| 55. | PDLIM3 | protein-protein interactions | DCTN1 |
| 56. | PDLIM3 | protein-protein interactions | FHL1 |
| 57. | PDLIM3 | protein-protein interactions | LMNA |
| 58. | PDLIM3 | protein-protein interactions | NAA40 |
| 59. | PDLIM3 | protein-protein interactions | PALLD |
| 60. | PDLIM3 | protein-protein interactions | PCNA |
| 61. | PDLIM3 | regulation of binding | ACTN2 |
| 62. | PMP22 | RNA-RNA interactions: non-targeting interactions | mir-9 (includes others) |
| 63. | PMP22 | protein-protein interactions | LMNA |
| 64. | PRPH | protein-protein interactions | MYC |
| 65. | PRPH | protein-protein interactions | MYCN |
| 66. | PRPH | protein-protein interactions | PDLIM3 |
| 67. | RANBP2 | protein-protein interactions | MYC |
| 68. | RANBP2 | protein-protein interactions | PDLIM3 |
| 69. | REST | protein-DNA interactions | mir-9 (includes others) |
| 70. | REST | protein-protein interactions | FHL1 |
| 71. | SQSTM1 | activation | AR |
| 72. | SQSTM1 | localization | AR |
| 73. | SQSTM1 | molecular cleavage | TARDBP |
| 74. | SQSTM1 | protein-protein interactions | AR |
| 75. | SQSTM1 | protein-protein interactions | FUS |

| | | | |
|---|---|---|---|
| 76. | SQSTM1 | protein-protein interactions | MYC |
| 77. | SQSTM1 | protein-protein interactions | PDLIM3 |
| 78. | SQSTM1 | protein-protein interactions | REST |
| 79. | SQSTM1 | protein-protein interactions | TARDBP |
| 80. | TARDBP | expression | mir-9 (includes others) |
| 81. | TARDBP | protein-protein interactions | AGR2 |
| 82. | TARDBP | protein-protein interactions | CUL3 |
| 83. | TARDBP | protein-protein interactions | NAA40 |
| 84. | TARDBP | protein-protein interactions | SQSTM1 |
| 85. | VIM | protein-protein interactions | AGO2 |
| 86. | VIM | protein-protein interactions | FUS |
| 87. | VIM | protein-protein interactions | FXR1 |
| 88. | VIM | protein-protein interactions | MYC |
| 89. | VIM | protein-protein interactions | MYCN |
| 90. | VIM | protein-protein interactions | PDLIM3 |
| 91. | YAP1 | protein-protein interactions | BAG3 |
| 92. | YAP1 | protein-protein interactions | LMNA |
| 93. | YAP1 | protein-protein interactions | PCNA |
| 94. | YAP1 | protein-protein interactions | PRPH |
| 95. | YAP1 | protein-protein interactions | VIM |
| 96. | YAP1 | transcription | mir-9 (includes others) |
| 97. | mir-9 (includes others) | RNA-RNA interactions: non-targeting interactions | AR |
| 98. | mir-9 (includes others) | protein-RNA interactions | AGO2 |
| 99. | mir-9 (includes others) | protein-RNA interactions | AGO3 |
| 100. | mir-9 (includes others) | protein-RNA interactions | FUS |
| 101. | mir-9 (includes others) | protein-RNA interactions | FXR1 |

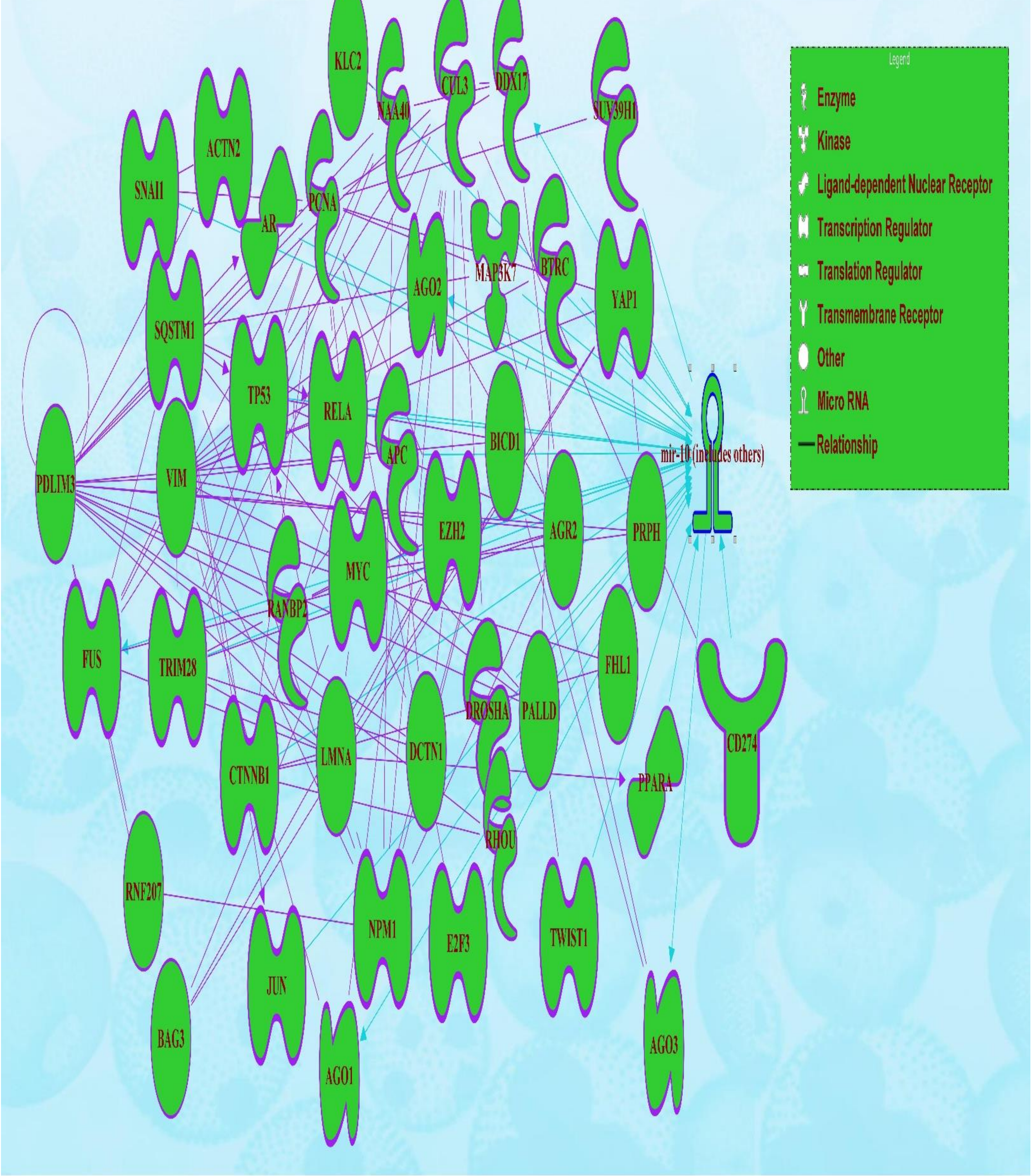


© 2000-2024 QIAGEN. All rights reserved.

Figure.8: Molecular network depicting the connectivity and relationships among the overlapping molecules associated with direct influence of *PDLIM-3* on miRNA-10 expression

**Table 11: Various molecules incorporated in the regulation of *PDLIM-3* and affect *miRNA10* expression**

| Symbol | Entrez Gene Name | Location | Family |
|---|---|---|---|
| 1. ACTN2 | actinin alpha 2 | Nucleus | transcription regulator |
| 2. AGO1 | argonaute RISC component 1 | Cytoplasm | translation regulator |
| 3. AGO2 | argonaute RISC catalytic component 2 | Cytoplasm | translation regulator |
| 4. AGO3 | argonaute RISC catalytic component 3 | Cytoplasm | translation regulator |
| 5. AGR2 | anterior gradient 2, protein disulphide isomerase family member | Extracellular Space | other |
| 6. APC | APC regulator of WNT signaling pathway | Nucleus | enzyme |
| 7. AR | androgen receptor | Nucleus | ligand-dependent nuclear receptor |
| 8. BAG3 | BAG cochaperone 3 | Cytoplasm | other |
| 9. BICD1 | BICD cargo adaptor 1 | Cytoplasm | other |
| 10. BTRC | beta-transducin repeat containing E3 ubiquitin protein ligase | Cytoplasm | enzyme |
| 11. CD274 | CD274 molecule | Plasma Membrane | transmembrane receptor |
| 12. CTNNB1 | catenin beta 1 | Nucleus | transcription regulator |
| 13. CUL3 | cullin 3 | Nucleus | enzyme |
| 14. DCTN1 | dynactin subunit 1 | Cytoplasm | other |
| 15. DDX17 | DEAD-box helicase 17 | Nucleus | enzyme |
| 16. DROSHA | drosha ribonuclease III | Nucleus | enzyme |
| 17. E2F3 | E2F transcription factor 3 | Nucleus | transcription regulator |
| 18. EZH2 | enhancer of zeste 2 polycomb repressive complex 2 subunit | Nucleus | transcription regulator |
| 19. FHL1 | four and a half LIM domains 1 | Cytoplasm | other |
| 20. FUS | FUS RNA binding protein | Nucleus | transcription regulator |
| 21. JUN | Jun proto-oncogene, AP-1 transcription factor subunit | Nucleus | transcription regulator |
| 22. KLC2 | kinesin light chain 2 | Cytoplasm | other |
| 23. LMNA | lamin A/C | Nucleus | other |
| 24. MAP3K7 | mitogen-activated protein kinase kinase kinase 7 | Cytoplasm | kinase |
| 25. mir-10 | relatives of microRNA 10 | Cytoplasm | microRNA |
| 26. MYC | MYC proto-oncogene, bHLH transcription factor | Nucleus | transcription regulator |
| 27. NAA40 | N-alpha-acetyltransferase 40, NatD catalytic subunit | Cytoplasm | enzyme |
| 28. NPM1 | nucleophosmin 1 | Nucleus | transcription regulator |
| 29. PALLD | palladin, cytoskeletal associated protein | Plasma Membrane | other |
| 30. PCNA | proliferating cell nuclear antigen | Nucleus | enzyme |
| 31. PDLIM3 | PDZ and LIM domain 3 | Cytoplasm | other |
| 32. PPARA | peroxisome proliferator activated receptor alpha | Nucleus | ligand-dependent nuclear receptor |
| 33. PRPH | peripherin | Plasma Membrane | other |
| 34. RANBP2 | RAN binding protein 2 | Nucleus | enzyme |
| 35. RELA | RELA proto-oncogene, NF-kB subunit | Nucleus | transcription regulator |
| 36. RHOU | ras homolog family member U | Cytoplasm | enzyme |
| 37. RNF207 | ring finger protein 207 | Other | other |
| 38. SNAI1 | snail family transcriptional repressor 1 | Nucleus | transcription regulator |

| | | | |
|---|---|---|---|
| 39. SQSTM1 | sequestosome 1 | Cytoplasm | transcription regulator |
| 40. SUV39H1 | SUV39H1 histone lysine methyltransferase | Nucleus | enzyme |
| 41. TP53 | tumor protein p53 | Nucleus | transcription regulator |
| 42. TRIM28 | tripartite motif containing 28 | Nucleus | transcription regulator |
| 43. TWIST1 | twist family bHLH transcription factor 1 | Nucleus | transcription regulator |
| 44. VIM | vimentin | Cytoplasm | other |
| 45. YAP1 | Yes1 associated transcriptional regulator | Nucleus | transcription regulator |

**Table.12: Various interactions between molecules incorporated in the *PDLIM-3* and affect *miRNA10* expression**

| From Molecule(s) | Relationship Type | To Molecule(s) |
|---|---|---|
| 1. ACTN2 | activation | AR |
| 2. ACTN2 | protein-protein interactions | AR |
| 3. AGR2 | inhibition | TP53 |
| 4. AGR2 | protein-protein interactions | AGO2 |
| 5. AGR2 | protein-protein interactions | TP53 |
| 6. AR | expression | mir-10 |
| 7. AR | protein-DNA interactions | mir-10 |
| 8. AR | protein-protein interactions | ACTN2 |
| 9. BICD1 | protein-protein interactions | AGO1 |
| 10. BICD1 | protein-protein interactions | AGO3 |
| 11. BTRC | protein-protein interactions | BICD1 |
| 12. CD274 | expression | mir-10 |
| 13. CTNNB1 | expression | mir-10 |
| 14. CUL3 | protein-protein interactions | AGO1 |
| 15. CUL3 | protein-protein interactions | AGO2 |
| 16. CUL3 | protein-protein interactions | AGO3 |
| 17. CUL3 | protein-protein interactions | CD274 |
| 18. CUL3 | protein-protein interactions | CTNNB1 |
| 19. DCTN1 | protein-protein interactions | APC |
| 20. DDX17 | protein-protein interactions | AGR2 |
| 21. DDX17 | protein-protein interactions | CUL3 |
| 22. DROSHA | protein-DNA interactions | mir-10 |
| 23. DROSHA | protein-protein interactions | CUL3 |
| 24. E2F3 | expression | mir-10 |
| 25. EZH2 | expression | mir-10 |
| 26. EZH2 | protein-protein interactions | BAG3 |
| 27. EZH2 | protein-protein interactions | CUL3 |
| 28. EZH2 | protein-protein interactions | DCTN1 |
| 29. FHL1 | protein-protein interactions | CTNNB1 |
| 30. FUS | protein-protein interactions | AGR2 |
| 31. FUS | protein-protein interactions | CUL3 |
| 32. LMNA | activation | TP53 |
| 33. LMNA | protein-DNA interactions | PPARA |
| 34. LMNA | protein-protein interactions | CTNNB1 |

| | | |
|---|---|---|
| 35. LMNA | protein-protein interactions | EZH2 |
| 36. LMNA | protein-protein interactions | FUS |
| 37. LMNA | protein-protein interactions | TP53 |
| 38. MYC | expression | mir-10 (includes others) |
| 39. MYC | protein-protein interactions | BAG3 |
| 40. MYC | protein-protein interactions | DCTN1 |
| 41. MYC | protein-protein interactions | LMNA |
| 42. NAA40 | protein-protein interactions | DDX17 |
| 43. NAA40 | protein-protein interactions | KLC2 |
| 44. NPM1 | expression | mir-10 (includes others) |
| 45. NPM1 | protein-protein interactions | AGR2 |
| 46. NPM1 | protein-protein interactions | BICD1 |
| 47. NPM1 | protein-protein interactions | CUL3 |
| 48. NPM1 | protein-protein interactions | LMNA |
| 49. NPM1 | protein-protein interactions | NAA40 |
| 50. PALLD | protein-protein interactions | BTRC |
| 51. PALLD | protein-protein interactions | MYC |
| 52. PCNA | protein-protein interactions | APC |
| 53. PCNA | protein-protein interactions | AR |
| 54. PCNA | protein-protein interactions | CTNNB1 |
| 55. PCNA | protein-protein interactions | DDX17 |
| 56. PCNA | protein-protein interactions | E2F3 |
| 57. PCNA | protein-protein interactions | EZH2 |
| 58. PCNA | protein-protein interactions | FUS |
| 59. PCNA | protein-protein interactions | MAP3K7 |
| 60. PCNA | protein-protein interactions | MYC |
| 61. PDLIM3 | protein-protein interactions | ACTN2 |
| 62. PDLIM3 | protein-protein interactions | AGR2 |
| 63. PDLIM3 | protein-protein interactions | BAG3 |
| 64. PDLIM3 | protein-protein interactions | BICD1 |
| 65. PDLIM3 | protein-protein interactions | CUL3 |
| 66. PDLIM3 | protein-protein interactions | DCTN1 |
| 67. PDLIM3 | protein-protein interactions | FHL1 |
| 68. PDLIM3 | protein-protein interactions | LMNA |
| 69. PDLIM3 | protein-protein interactions | NAA40 |
| 70. PDLIM3 | protein-protein interactions | PALLD |
| 71. PDLIM3 | protein-protein interactions | PCNA |
| 72. PDLIM3 | protein-protein interactions | PDLIM3 |
| 73. PPARA | expression | mir-10 |
| 74. PRPH | protein-protein interactions | MYC |
| 75. PRPH | protein-protein interactions | PDLIM3 |
| 76. RANBP2 | protein-protein interactions | APC |
| 77. RANBP2 | protein-protein interactions | CTNNB1 |

| | | |
|---|---|---|
| 78. RANBP2 | protein-protein interactions | EZH2 |
| 79. RANBP2 | protein-protein interactions | MYC |
| 80. RANBP2 | protein-protein interactions | PDLIM3 |
| 81. RELA | expression | mir-10 (includes others) |
| 82. RELA | protein-DNA interactions | mir-10 (includes others) |
| 83. RELA | protein-protein interactions | PCNA |
| 84. RHOU | protein-protein interactions | CTNNB1 |
| 85. RHOU | protein-protein interactions | PDLIM3 |
| 86. RNF207 | protein-protein interactions | NPM1 |
| 87. RNF207 | protein-protein interactions | PDLIM3 |
| 88. SNAI1 | expression | mir-10 (includes others) |
| 89. SNAI1 | protein-protein interactions | ACTN2 |
| 90. SNAI1 | protein-protein interactions | PCNA |
| 91. SNAI1 | transcription | mir-10 (includes others) |
| 92. SQSTM1 | activation | AR |
| 93. SQSTM1 | activation | TP53 |
| 94. SQSTM1 | expression | RELA |
| 95. SQSTM1 | expression | SNAI1 |
| 96. SQSTM1 | protein-protein interactions | AR |
| 97. SQSTM1 | protein-protein interactions | CTNNB1 |
| 98. SQSTM1 | protein-protein interactions | FUS |
| 99. SQSTM1 | protein-protein interactions | JUN |
| 100. SQSTM1 | protein-protein interactions | MAP3K7 |
| 101. SQSTM1 | protein-protein interactions | MYC |
| 102. SQSTM1 | protein-protein interactions | NPM1 |
| 103. SQSTM1 | protein-protein interactions | PDLIM3 |
| 104. SQSTM1 | protein-protein interactions | RELA |
| 105. SQSTM1 | protein-protein interactions | SNAI1 |
| 106. SQSTM1 | protein-protein interactions | TP53 |
| 107. SQSTM1 | transcription | JUN |
| 108. SUV39H1 | protein-protein interactions | PCNA |
| 109. SUV39H1 | protein-protein interactions | mir-10 |
| 110. TP53 | expression | mir-10 |
| 111. TP53 | protein-protein interactions | AGR2 |
| 112. TP53 | protein-protein interactions | BICD1 |
| 113. TP53 | protein-protein interactions | CUL3 |
| 114. TP53 | protein-protein interactions | LMNA |
| 115. TP53 | protein-protein interactions | NAA40 |
| 116. TP53 | protein-protein interactions | PALLD |
| 117. TP53 | protein-protein interactions | PCNA |
| 118. TP53 | protein-protein interactions | RANBP2 |

| | | |
|---|---|---|
| 119. TP53 | protein-protein interactions | SQSTM1 |
| 120. TRIM28 | expression | mir-10 |
| 121. TRIM28 | protein-protein interactions | AGR2 |
| 122. TRIM28 | protein-protein interactions | CUL3 |
| 123. TRIM28 | protein-protein interactions | DCTN1 |
| 124. TRIM28 | protein-protein interactions | LMNA |
| 125. TRIM28 | protein-protein interactions | NAA40 |
| 126. TRIM28 | protein-protein interactions | PCNA |
| 127. TRIM28 | protein-protein interactions | RANBP2 |
| 128. TWIST1 | expression | mir-10 |
| 129. TWIST1 | protein-protein interactions | PALLD |
| 130. VIM | protein-protein interactions | AGO1 |
| 131. VIM | protein-protein interactions | AGO2 |
| 132. VIM | protein-protein interactions | APC |
| 133. VIM | protein-protein interactions | BTRC |
| 134. VIM | protein-protein interactions | EZH2 |
| 135. VIM | protein-protein interactions | FUS |
| 136. VIM | protein-protein interactions | MYC |
| 137. VIM | protein-protein interactions | NPM1 |
| 138. VIM | protein-protein interactions | PDLIM3 |
| 139. VIM | protein-protein interactions | RELA |
| 140. VIM | protein-protein interactions | TP53 |
| 141. VIM | protein-protein interactions | TRIM28 |
| 142. YAP1 | protein-protein interactions | BAG3 |
| 143. YAP1 | protein-protein interactions | LMNA |
| 144. YAP1 | protein-protein interactions | PCNA |
| 145. YAP1 | protein-protein interactions | PRPH |
| 146. YAP1 | protein-protein interactions | VIM |
| 147. YAP1 | transcription | mir-10 |
| 148. mir-10 | RNA-RNA interactions: non-targeting interactions | APC |
| 149. mir-10 | RNA-RNA interactions: non-targeting interactions | BTRC |
| 150. mir-10 | RNA-RNA interactions: non-targeting interactions | JUN |
| 151. mir-10 | RNA-RNA interactions: non-targeting interactions | MAP3K7 |
| 152. mir-10 | protein-RNA interactions | AGO1 |
| 153. mir-10 | protein-RNA interactions | AGO2 |
| 154. mir-10 | protein-RNA interactions | AGO3 |
| 155. mir-10 | protein-RNA interactions | DDX17 |
| 156. mir-10 | protein-RNA interactions | FUS |
| 157. mir-10 | protein-protein interactions | KLC2 |

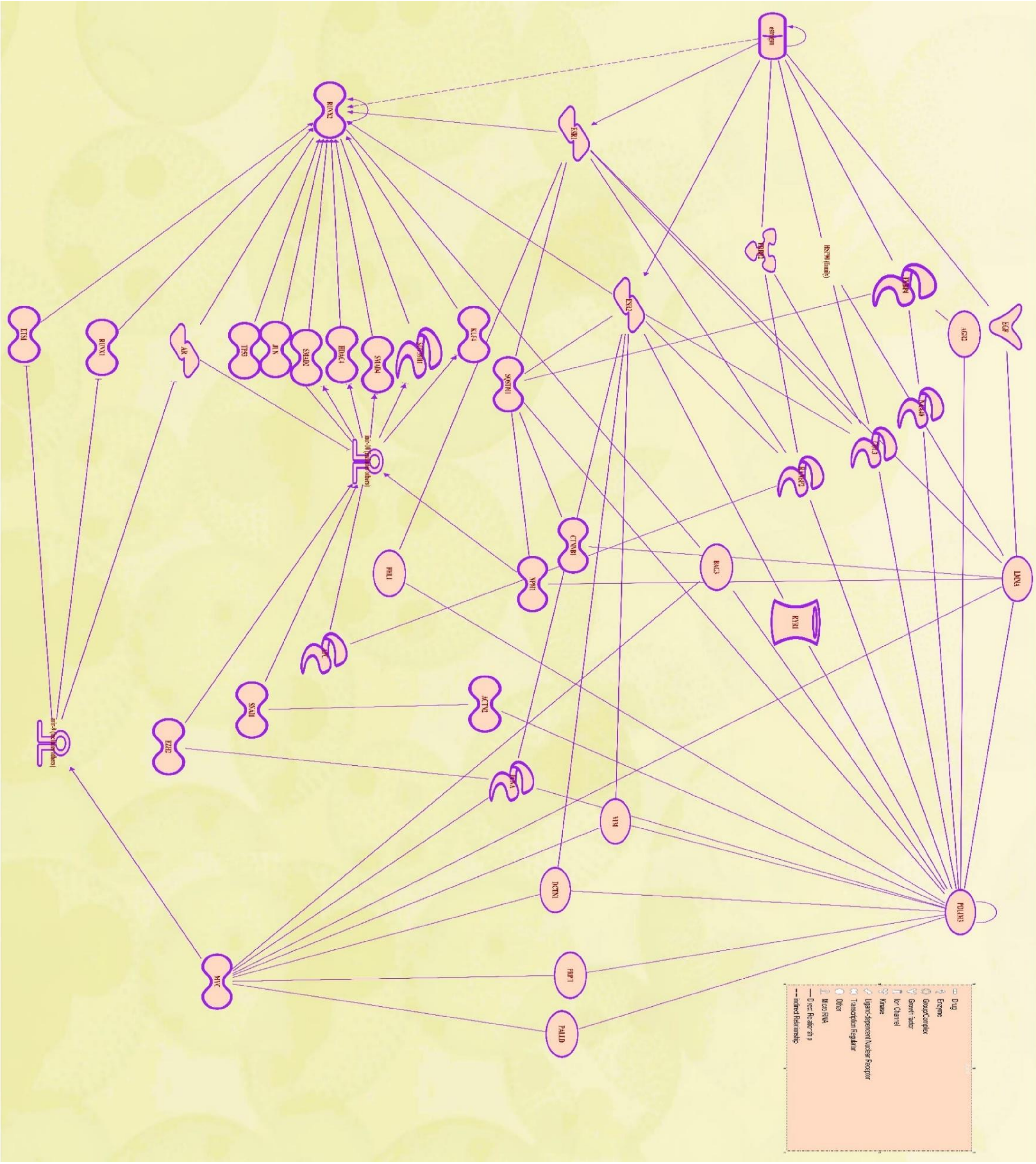

**Figure.9: Molecules network influenced by estrogen, *PDLIM-3*. miRNA-10, miRNA-9 affecting RUNX-2expression**

**Table 13: Molecules regulated by Estrogen, PDLIM-3, miRNA9 and miRNA10 affecting RUNX-2 expression**

| Symbol | Gene Name | Location | Family |
|---|---|---|---|
| 1. ACTN2 | actinin alpha 2 | Nucleus | transcription regulator |
| 2. AGR2 | anterior gradient 2, protein disulphide isomerase family member | Extracellular Space | other |
| 3. APC | APC regulator of WNT signaling pathway | Nucleus | enzyme |
| 4. AR | androgen receptor | Nucleus | ligand-dependent nuclear receptor |
| 5. BAG3 | BAG cochaperone 3 | Cytoplasm | Other |
| 6. CTNNB1 | catenin beta 1 | Nucleus | transcription regulator |
| 7. CUL3 | cullin 3 | Nucleus | Enzyme |
| 8. DCTN1 | dynactin subunit 1 | Cytoplasm | Other |
| 9. EGF | epidermal growth factor | Extracellular Space | growth factor |
| 10. ERBB2 | erb-b2 receptor tyrosine kinase 2 | Plasma Membrane | Kinase |
| 11. ESR1 | estrogen receptor 1 | Nucleus | ligand-dependent nuclear receptor |
| 12. ESR2 | estrogen receptor 2 | Nucleus | ligand-dependent nuclear receptor |
| 13. ETS1 | ETS proto-oncogene 1, transcription factor | Nucleus | transcription regulator |
| 14. EZH2 | enhancer of zeste 2 polycomb repressive complex 2 subunit | Nucleus | transcription regulator |
| 15. FHL1 | four and a half LIM domains 1 | Cytoplasm | other |
| 16. FKBP4 | FKBP prolyl isomerase 4 | Nucleus | enzyme |
| 17. HDAC4 | histone deacetylase 4 | Nucleus | transcription regulator |
| 18. HSP90 (family) | | Cytoplasm | group |
| 19. JUN | Jun proto-oncogene, AP-1 transcription factor subunit | Nucleus | transcription regulator |
| 20. KLF4 | KLF transcription factor 4 | Nucleus | transcription regulator |
| 21. LMNA | lamin A/C | Nucleus | other |
| 22. mir-10 | relatives of microRNA 10 | Cytoplasm | microRNA |
| 23. mir-9 | relatives of microRNA 9 | Cytoplasm | microRNA |
| 24. MYC | MYC proto-oncogene, bHLH transcription factor | Nucleus | transcription regulator |
| 25. NAA40 | N-alpha-acetyltransferase 40, NatD catalytic subunit | Cytoplasm | enzyme |
| 26. NPM1 | nucleophosmin 1 | Nucleus | transcription regulator |

| | | | |
|---|---|---|---|
| 27. PALLD | palladin, cytoskeletal associated protein | Plasma Membrane | other |
| 28. PCNA | proliferating cell nuclear antigen | Nucleus | enzyme |
| 29. PDLIM3 | PDZ and LIM domain 3 | Cytoplasm | other |
| 30. PRPH | peripherin | Plasma Membrane | other |
| 31. RANBP2 | RAN binding protein 2 | Nucleus | enzyme |
| 32. RUNX1 | RUNX family transcription factor 1 | Nucleus | transcription regulator |
| 33. RUNX2 | RUNX family transcription factor 2 | Nucleus | transcription regulator |
| 34. RYR1 | ryanodine receptor 1 | Cytoplasm | ion channel |
| 35. SMAD2 | SMAD family member 2 | Nucleus | transcription regulator |
| 36. SMAD4 | SMAD family member 4 | Nucleus | transcription regulator |
| 37. SNAI1 | snail family transcriptional repressor 1 | Nucleus | transcription regulator |
| 38. SQSTM1 | sequestosome 1 | Cytoplasm | transcription regulator |
| 39. SUV39H1 | SUV39H1 histone lysine methyltransferase | Nucleus | enzyme |
| 40. TP53 | tumor protein p53 | Nucleus | transcription regulator |
| 41. VIM | vimentin | Cytoplasm | other |

**Table.14: Various interactions regulated by Estrogen, PDLIM-3, miRNA9 and miRNA10 affecting RUNX-2**

| From Molecule(s) | Relationship Type | To Molecule(s) |
|---|---|---|
| 1. AR | expression | mir-10 (includes others) |
| 2. AR | protein-DNA interactions | mir-10 (includes others) |
| 3. ERBB2 | protein-protein interactions | CUL3 |
| 4. ESR1 | activation | RUNX2 |
| 5. ESR1 | chemical-protein interactions | estrogen |
| 6. ESR1 | expression | RUNX2 |
| 7. ESR1 | inhibition | RUNX2 |
| 8. ESR1 | protein-protein interactions | CUL3 |
| 9. ESR1 | protein-protein interactions | RUNX2 |
| 10. ESR1 | transcription | RUNX2 |
| 11. ESR2 | chemical-protein interactions | estrogen |
| 12. ESR2 | expression | RUNX2 |
| 13. ESR2 | protein-protein interactions | CUL3 |
| 14. ESR2 | protein-protein interactions | DCTN1 |
| 15. ETS1 | protein-DNA interactions | RUNX2 |
| 16. ETS1 | protein-protein interactions | RUNX2 |
| 17. ETS1 | transcription | RUNX2 |
| 18. EZH2 | expression | mir-10 (includes others) |
| 19. FHL1 | protein-protein interactions | ESR1 |
| 20. FKBP4 | chemical-protein interactions | estrogen |
| 21. FKBP4 | protein-protein interactions | AGR2 |
| 22. HDAC4 | activation | RUNX2 |

| | | |
|---|---|---|
| 23. HDAC4 | expression | RUNX2 |
| 24. HDAC4 | protein-protein interactions | RUNX2 |
| 25. HSP90 (family) | chemical-protein interactions | estrogen |
| 26. HSP90 (family) | protein-protein interactions | CUL3 |
| 27. JUN | protein-DNA interactions | RUNX2 |
| 28. JUN | protein-protein interactions | RUNX2 |
| 29. KLF4 | expression | RUNX2 |
| 30. KLF4 | protein-DNA interactions | RUNX2 |
| 31. KLF4 | protein-protein interactions | RUNX2 |
| 32. LMNA | protein-protein interactions | CTNNB1 |
| 33. LMNA | protein-protein interactions | EGF |
| 34. LMNA | protein-protein interactions | ESR1 |
| 35. LMNA | protein-protein interactions | HSP90 (family) |
| 36. MYC | expression | mir-9 (includes others) |
| 37. MYC | protein-DNA interactions | mir-9 (includes others) |
| 38. MYC | protein-protein interactions | BAG3 |
| 39. MYC | protein-protein interactions | DCTN1 |
| 40. MYC | protein-protein interactions | LMNA |
| 41. MYC | transcription | mir-9 (includes others) |
| 42. NAA40 | protein-protein interactions | FKBP4 |
| 43. NPM1 | expression | mir-10 (includes others) |
| 44. NPM1 | protein-protein interactions | LMNA |
| 45. PALLD | protein-protein interactions | MYC |
| 46. PCNA | protein-protein interactions | ESR2 |
| 47. PCNA | protein-protein interactions | EZH2 |
| 48. PCNA | protein-protein interactions | MYC |
| 49. PDLIM3 | protein-protein interactions | ACTN2 |
| 50. PDLIM3 | protein-protein interactions | AGR2 |
| 51. PDLIM3 | protein-protein interactions | BAG3 |
| 52. PDLIM3 | protein-protein interactions | CUL3 |
| 53. PDLIM3 | protein-protein interactions | DCTN1 |
| 54. PDLIM3 | protein-protein interactions | FHL1 |
| 55. PDLIM3 | protein-protein interactions | LMNA |
| 56. PDLIM3 | protein-protein interactions | NAA40 |
| 57. PDLIM3 | protein-protein interactions | PALLD |
| 58. PDLIM3 | protein-protein interactions | PCNA |
| 59. PDLIM3 | protein-protein interactions | PDLIM3 |
| 60. PRPH | protein-protein interactions | MYC |
| 61. PRPH | protein-protein interactions | PDLIM3 |
| 62. RANBP2 | protein-protein interactions | APC |
| 63. RANBP2 | protein-protein interactions | ERBB2 |
| 64. RANBP2 | protein-protein interactions | ESR1 |
| 65. RANBP2 | protein-protein interactions | ESR2 |
| 66. RANBP2 | protein-protein interactions | PDLIM3 |

| | | |
|---|---|---|
| 67. RUNX1 | expression | RUNX2 |
| 68. RUNX1 | protein-DNA interactions | RUNX2 |
| 69. RUNX1 | protein-protein interactions | RUNX2 |
| 70. RUNX2 | activation | RUNX2 |
| 71. RUNX2 | expression | RUNX2 |
| 72. RUNX2 | inhibition | RUNX2 |
| 73. RUNX2 | localization | RUNX2 |
| 74. RUNX2 | modification | RUNX2 |
| 75. RUNX2 | molecular cleavage | RUNX2 |
| 76. RUNX2 | protein-DNA interactions | RUNX2 |
| 77. RUNX2 | protein-protein interactions | AR |
| 78. RUNX2 | protein-protein interactions | BAG3 |
| 79. RUNX2 | protein-protein interactions | ESR1 |
| 80. RUNX2 | protein-protein interactions | ETS1 |
| 81. RUNX2 | protein-protein interactions | HDAC4 |
| 82. RUNX2 | protein-protein interactions | JUN |
| 83. RUNX2 | protein-protein interactions | KLF4 |
| 84. RUNX2 | protein-protein interactions | RUNX1 |
| 85. RUNX2 | protein-protein interactions | SMAD4 |
| 86. RUNX2 | regulation of binding | RUNX2 |
| 87. RUNX2 | ubiquitination | RUNX2 |
| 88. RYR1 | protein-protein interactions | ESR2 |
| 89. RYR1 | protein-protein interactions | PDLIM3 |
| 90. SMAD2 | expression | RUNX2 |
| 91. SMAD2 | protein-protein interactions | RUNX2 |
| 92. SMAD4 | expression | RUNX2 |
| 93. SMAD4 | protein-DNA interactions | RUNX2 |
| 94. SMAD4 | protein-protein interactions | RUNX2 |
| 95. SNAI1 | expression | mir-10 (includes others) |
| 96. SNAI1 | protein-protein interactions | ACTN2 |
| 97. SNAI1 | transcription | mir-10 (includes others) |
| 98. SQSTM1 | protein-protein interactions | CTNNB1 |
| 99. SQSTM1 | protein-protein interactions | ESR1 |
| 100. SQSTM1 | protein-protein interactions | ESR2 |
| 101. SQSTM1 | protein-protein interactions | FKBP4 |
| 102. SQSTM1 | protein-protein interactions | NPM1 |
| 103. SQSTM1 | protein-protein interactions | PDLIM3 |
| 104. SUV39H1 | protein-protein interactions | RUNX2 |
| 105. SUV39H1 | protein-protein interactions | mir-10 (includes others) |
| 106. TP53 | expression | RUNX2 |
| 107. TP53 | protein-protein interactions | RUNX2 |
| 108. TP53 | transcription | RUNX2 |
| 109. VIM | protein-protein interactions | ESR2 |
| 110. VIM | protein-protein interactions | MYC |

| | | |
|---|---|---|
| 111. VIM | protein-protein interactions | PDLIM3 |
| 112. estrogen | activation | ESR1 |
| 113. estrogen | activation | ESR2 |
| 114. estrogen | activation | RUNX2 |
| 115. estrogen | chemical-protein interactions | EGF |
| 116. estrogen | chemical-protein interactions | ERBB2 |
| 117. estrogen | chemical-protein interactions | ESR1 |
| 118. estrogen | chemical-protein interactions | ESR2 |
| 119. estrogen | translocation | estrogen |
| 120. mir-10 (includes others) | RNA-RNA interactions: microRNA targeting | AR |
| 121. mir-10 (includes others) | RNA-RNA interactions: microRNA targeting | HDAC4 |
| 122. mir-10 (includes others) | RNA-RNA interactions: microRNA targeting | KLF4 |
| 123. mir-10 (includes others) | RNA-RNA interactions: microRNA targeting | SMAD2 |
| 124. mir-10 (includes others) | RNA-RNA interactions: microRNA targeting | SMAD4 |
| 125. mir-10 (includes others) | RNA-RNA interactions: microRNA targeting | SUV39H1 |
| 126. mir-10 (includes others) | RNA-RNA interactions: non-targeting interactions | APC |
| 127. mir-10 (includes others) | RNA-RNA interactions: non-targeting interactions | JUN |
| 128. mir-10 (includes others) | expression | AR |
| 129. mir-10 (includes others) | protein-protein interactions | SUV39H1 |
| 130. mir-9 (includes others) | RNA-RNA interactions: microRNA targeting | AR |
| 131. mir-9 (includes others) | RNA-RNA interactions: microRNA targeting | ETS1 |
| 132. mir-9 (includes others) | RNA-RNA interactions: microRNA targeting | RUNX1 |
| 133. mir-9 (includes others) | RNA-RNA interactions: non-targeting interactions | AR |
| 134. mir-9 (includes others) | expression | AR |
| 135. mir-9 (includes others) | expression | ETS1 |
| 136. mir-9 (includes others) | expression | RUNX1 |

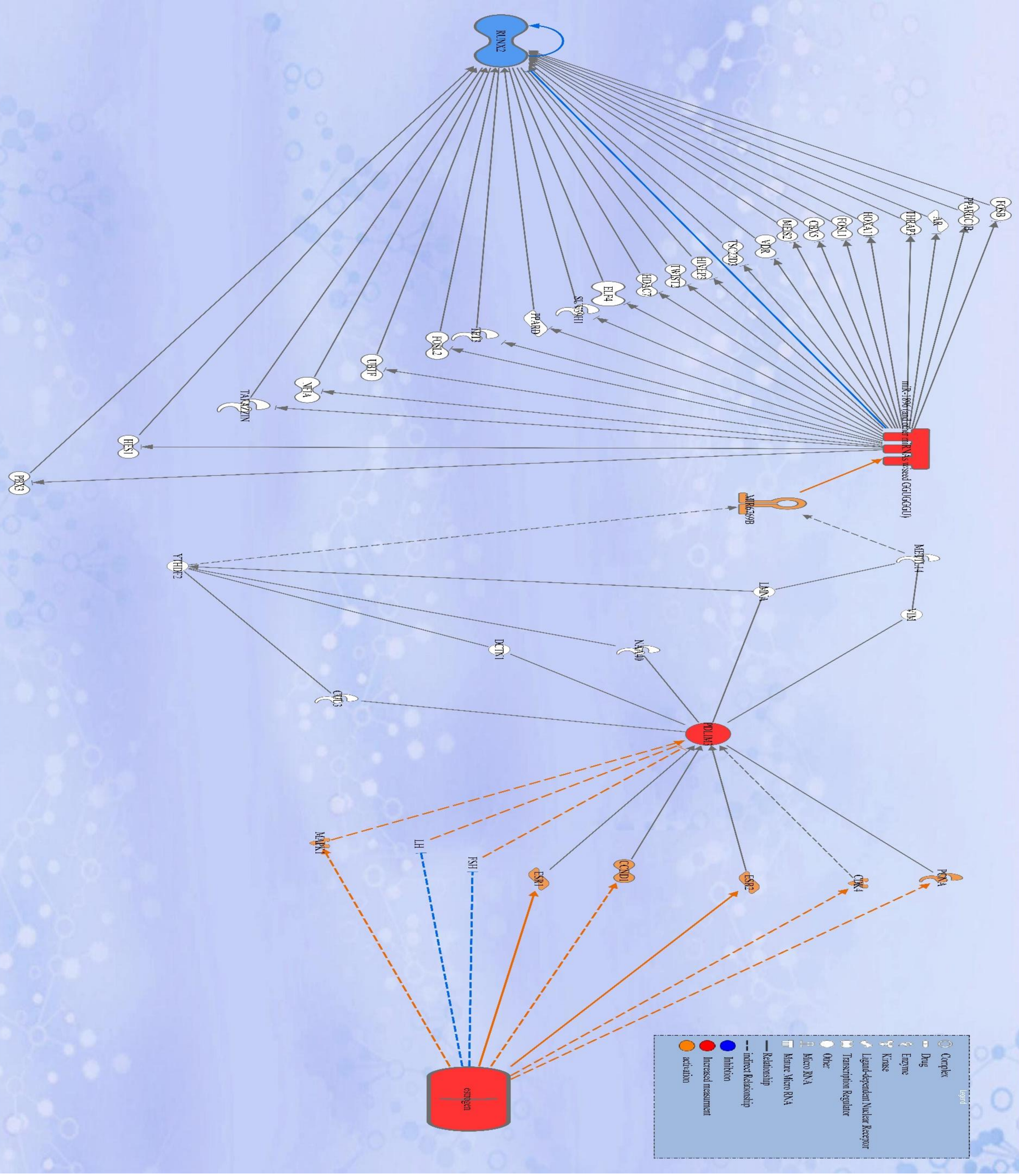


**Figure.10: Molecules network regulated by estrogen, *PDLIM-3*. microRNA-1896 , microRNA6769B affecting RUNX-2expression**

Table 15: Molecules regulated by Estrogen, PDLIM-3, , microRNA-1896 , microRNA6769B, affecting RUNX-2

| Symbol | Molecule/ Gene Name | Location | Family |
|---|---|---|---|
| 1. AR | androgen receptor | Nucleus | ligand-dependent nuclear receptor |
| 2. CBX5 | chromobox 5 | Nucleus | transcription regulator |
| 3. CCND1 | cyclin D1 | Nucleus | transcription regulator |
| 4. CDK4 | cyclin dependent kinase 4 | Nucleus | kinase |
| 5. CUL3 | cullin 3 | Nucleus | Enzyme |
| 6. DCTN1 | dynactin subunit 1 | Cytoplasm | Other |
| 7. ELF4 | E74 like ETS transcription factor 4 | Nucleus | transcription regulator |
| 8. ESR1 | estrogen receptor 1 | Nucleus | ligand-dependent nuclear receptor |
| 9. ESR2 | estrogen receptor 2 | Nucleus | ligand-dependent nuclear receptor |
| 10. estrogen | | Other | chemical drug |
| 11. FOSB | FosB proto-oncogene, AP-1 transcription factor subunit | Nucleus | transcription regulator |
| 12. FOSL1 | FOS like 1, AP-1 transcription factor subunit | Nucleus | transcription regulator |
| 13. FOSL2 | FOS like 2, AP-1 transcription factor subunit | Nucleus | transcription regulator |
| 14. FSH | | Plasma Membrane | Complex |
| 15. HDAC7 | histone deacetylase 7 | Nucleus | transcription regulator |
| 16. HES1 | hes family bHLH transcription factor 1 | Nucleus | transcription regulator |
| 17. HIVEP3 | HIVEP zinc finger 3 | Nucleus | transcription regulator |
| 18. HOXA11 | homeobox A11 | Nucleus | transcription regulator |
| 19. LH | | Plasma Membrane | Complex |
| 20. LMNA | lamin A/C | Nucleus | Other |
| 21. MAPK1 | mitogen-activated protein kinase 1 | Cytoplasm | Kinase |
| 22. MEIS2 | Meis homeobox 2 | Nucleus | transcription regulator |
| 23. METTL14 | methyltransferase 14, N6-adenosine-methyltransferase subunit | Nucleus | Enzyme |
| 24. miR-1896 | | Cytoplasm | mature microRNA |
| 25. MIR6769B | microRNA 6769b | Other | microRNA |
| 26. NAA40 | N-alpha-acetyltransferase 40, NatD catalytic subunit | Cytoplasm | Enzyme |
| 27. NFIA | nuclear factor I A | Nucleus | transcription regulator |
| 28. PBX3 | PBX homeobox 3 | Nucleus | transcription regulator |
| 29. PCNA | proliferating cell nuclear antigen | Nucleus | Enzyme |
| 30. PDLIM3 | PDZ and LIM domain 3 | Cytoplasm | Other |
| 31. PPARD | peroxisome proliferator activated receptor delta | Nucleus | ligand-dependent nuclear receptor |
| 32. PPARGC1B | PPARG coactivator 1 beta | Nucleus | transcription regulator |
| 33. RUNX2 | RUNX family transcription factor 2 | Nucleus | transcription regulator |
| 34. SUV39H1 | SUV39H1 histone lysine methyltransferase | Nucleus | Enzyme |
| 35. TAFAZZIN | tafazzin, phospholipid-lysophospholipid transacylase | Nucleus | Enzyme |
| 36. TET3 | tet methylcytosine dioxygenase 3 | Nucleus | Enzyme |
| 37. THRAP3 | thyroid hormone receptor associated protein 3 | Nucleus | transcription regulator |
| 38. TSC22D3 | TSC22 domain family member 3 | Nucleus | transcription regulator |
| 39. TWIST2 | twist family bHLH transcription factor 2 | Nucleus | transcription regulator |
| 40. UBTF | upstream binding transcription factor | Nucleus | transcription regulator |
| 41. VDR | vitamin D receptor | Nucleus | transcription regulator |
| 42. VIM | vimentin | Cytoplasm | Other |
| 43. YTHDF2 | YTH N6-methyladenosine RNA binding protein F2 | Cytoplasm | Other |

**Table 16: Various interactions regulated by estrogen, *PDLIM-3*. microRNA-1896 , microRNA6769B affecting RUNX-2**

| From Molecule(s) | Relationship Type | To Molecule(s) |
|---|---|---|
| 1. CBX5 | expression | RUNX2 |
| 2. CCND1 | expression | PDLIM3 |

| | | |
|---|---|---|
| 3. CDK4 | expression | PDLIM3 |
| 4. ESR1 | chemical-protein interactions | estrogen |
| 5. ESR1 | expression | PDLIM3 |
| 6. ESR1 | regulation of binding | estrogen |
| 7. ESR2 | chemical-protein interactions | estrogen |
| 8. ESR2 | expression | PDLIM3 |
| 9. FOSB | protein-DNA interactions | RUNX2 |
| 10. FOSL1 | protein-DNA interactions | RUNX2 |
| 11. FOSL2 | protein-DNA interactions | RUNX2 |
| 12. FSH | expression | PDLIM3 |
| 13. HES1 | activation | RUNX2 |
| 14. HES1 | protein-protein interactions | RUNX2 |
| 15. HIVEP3 | inhibition | RUNX2 |
| 16. HIVEP3 | protein-protein interactions | RUNX2 |
| 17. HOXA11 | expression | RUNX2 |
| 18. HOXA11 | protein-protein interactions | RUNX2 |
| 19. HOXA11 | transcription | RUNX2 |
| 20. LH | expression | PDLIM3 |
| 21. LMNA | protein-DNA interactions | YTHDF2 |
| 22. MAPK1 | expression | PDLIM3 |
| 23. MEIS2 | expression | RUNX2 |
| 24. METTL14 | expression | MIR6769B |
| 25. METTL14 | protein-protein interactions | LMNA |
| 26. MIR6769B | processing yields | miR-1896 |
| 27. NFIA | expression | RUNX2 |
| 28. PBX3 | expression | RUNX2 |
| 29. PDLIM3 | protein-protein interactions | CUL3 |
| 30. PDLIM3 | protein-protein interactions | DCTN1 |
| 31. PDLIM3 | protein-protein interactions | LMNA |
| 32. PDLIM3 | protein-protein interactions | NAA40 |
| 33. PDLIM3 | protein-protein interactions | PCNA |
| 34. PPARD | expression | RUNX2 |
| 35. PPARGC1B | transcription | RUNX2 |
| 36. RUNX2 | activation | RUNX2 |
| 37. RUNX2 | expression | RUNX2 |
| 38. RUNX2 | inhibition | HIVEP3 |
| 39. RUNX2 | inhibition | RUNX2 |
| 40. RUNX2 | localization | RUNX2 |
| 41. RUNX2 | modification | RUNX2 |
| 42. RUNX2 | molecular cleavage | RUNX2 |
| 43. RUNX2 | protein-DNA interactions | RUNX2 |
| 44. RUNX2 | protein-protein interactions | AR |
| 45. RUNX2 | protein-protein interactions | ELF4 |
| 46. RUNX2 | protein-protein interactions | HDAC7 |

| | | |
|---|---|---|
| 47. RUNX2 | protein-protein interactions | HES1 |
| 48. RUNX2 | protein-protein interactions | HIVEP3 |
| 49. RUNX2 | protein-protein interactions | HOXA11 |
| 50. RUNX2 | protein-protein interactions | TAFAZZIN |
| 51. RUNX2 | regulation of binding | RUNX2 |
| 52. RUNX2 | ubiquitination | RUNX2 |
| 53. SUV39H1 | protein-protein interactions | RUNX2 |
| 54. TAFAZZIN | protein-protein interactions | RUNX2 |
| 55. TET3 | protein-protein interactions | RUNX2 |
| 56. THRAP3 | expression | RUNX2 |
| 57. TSC22D3 | expression | RUNX2 |
| 58. TWIST2 | protein-protein interactions | RUNX2 |
| 59. UBTF | protein-protein interactions | RUNX2 |
| 60. VDR | expression | RUNX2 |
| 61. VDR | protein-protein interactions | RUNX2 |
| 62. VIM | protein-protein interactions | METTL14 |
| 63. VIM | protein-protein interactions | PDLIM3 |
| 64. YTHDF2 | expression | MIR6769B |
| 65. YTHDF2 | protein-protein interactions | CUL3 |
| 66. YTHDF2 | protein-protein interactions | DCTN1 |
| 67. YTHDF2 | protein-protein interactions | NAA40 |
| 68. estrogen | activation | CDK4 |
| 69. estrogen | activation | ESR1 |
| 70. estrogen | activation | ESR2 |
| 71. estrogen | activation | MAPK1 |
| 72. estrogen | chemical-protein interactions | ESR1 |
| 73. estrogen | chemical-protein interactions | ESR2 |
| 74. estrogen | expression | CCND1 |
| 75. estrogen | expression | ESR1 |
| 76. estrogen | expression | ESR2 |
| 77. estrogen | expression | FSH |
| 78. estrogen | expression | LH |
| 79. estrogen | expression | PCNA |
| 80. estrogen | localization | FSH |
| 81. estrogen | localization | LH |
| 82. estrogen | molecular cleavage | ESR1 |
| 83. estrogen | phosphorylation | ESR1 |
| 84. estrogen | phosphorylation | ESR2 |
| 85. estrogen | phosphorylation | MAPK1 |
| 86. estrogen | regulation of binding | CCND1 |
| 87. estrogen | regulation of binding | CDK4 |
| 88. estrogen | regulation of binding | ESR1 |
| 89. estrogen | regulation of binding | ESR2 |
| 90. estrogen | transcription | CCND1 |

| | | |
|---|---|---|
| 91. estrogen | translocation | ESR1 |
| 92. estrogen | translocation | ESR2 |
| 93. miR-1896 | RNA-RNA interactions: microRNA targeting | AR |
| 94. miR-1896 | RNA-RNA interactions: microRNA targeting | CBX5 |
| 95. miR-1896 | RNA-RNA interactions: microRNA targeting | ELF4 |
| 96. miR-1896 | RNA-RNA interactions: microRNA targeting | FOSB |
| 97. miR-1896 | RNA-RNA interactions: microRNA targeting | FOSL1 |
| 98. miR-1896 | RNA-RNA interactions: microRNA targeting | FOSL2 |
| 99. miR-1896 | RNA-RNA interactions: microRNA targeting | HDAC7 |
| 100. miR-1896 | RNA-RNA interactions: microRNA targeting | HES1 |
| 101. miR-1896 | RNA-RNA interactions: microRNA targeting | HIVEP3 |
| 102. miR-1896 | RNA-RNA interactions: microRNA targeting | HOXA11 |
| 103. miR-1896 | RNA-RNA interactions: microRNA targeting | MEIS2 |
| 104. miR-1896 | RNA-RNA interactions: microRNA targeting | NFIA |
| 105. miR-1896 | RNA-RNA interactions: microRNA targeting | PBX3 |
| 106. miR-1896 | RNA-RNA interactions: microRNA targeting | PPARD |
| 107. miR-1896 | RNA-RNA interactions: microRNA targeting | PPARGC1B |
| 108. miR-1896 | RNA-RNA interactions: microRNA targeting | RUNX2 |
| 109. miR-1896 | RNA-RNA interactions: microRNA targeting | SUV39H1 |
| 110. miR-1896 | RNA-RNA interactions: microRNA targeting | TAFAZZIN |
| 111. miR-1896 | RNA-RNA interactions: microRNA targeting | TET3 |
| 112. miR-1896 | RNA-RNA interactions: microRNA targeting | THRAP3 |
| 113. miR-1896 | RNA-RNA interactions: microRNA targeting | TSC22D3 |
| 114. miR-1896 | RNA-RNA interactions: microRNA targeting | TWIST2 |
| 115. miR-1896 | RNA-RNA interactions: microRNA targeting | UBTF |
| 116. miR-1896 | RNA-RNA interactions: microRNA targeting | VDR |

| | | | |
|---|---|---|---|
| 117. | miR-1896 | expression | RUNX2 |
